# Embryonic Mast Cells Arise from *Cpa3*-expressing Precursors Independent of Granulocyte-Monocyte Progenitors

**DOI:** 10.1101/2024.10.30.620640

**Authors:** Wei Ma, Huifang Chen, Fei Gao, Han Zhao, Ningbo Wu, Shuangyan Zhang, Yiwen Zhu, Zijian Xu, Yu Lan, Bing Liu, Youqiong Ye, Zhaoyuan Liu, Florent Ginhoux, Bing Su

## Abstract

Most of the mast cells (MCs) in connective tissues, such as skin, are long-lived embryonic-derived immune cells that play important roles in host defense and various immunological diseases, including allergies. Their embryonic origin and ontogeny remain to be fully studied since several overlapping waves of embryonic hematopoiesis have been shown to give rise to these cells. Here, combining single-cell RNA sequencing and new genetic fate mapping models, we identified a *Cpa3*-expressing population sequentially appearing in the yolk sac, fetal liver, and peripheral tissues which gives rise to dermal MCs during embryonic days 11.5 to 14.5. Using in vitro differentiation and in vivo transplant methods, we identified a Ter119^−^F4/80^−^ CD45^+^CD117^+^CD16/32^+^CD135^−^CD115^−^Ly6C^−^CD34^lo^ population as potential fetal liver MC precursors. Fate mapping with *Cpa3^CreERT2^*and *Zbtb16^CreERT2^* models, as well as the granulocyte-monocyte progenitor (GMP) models *Ms4a3^Cre^* and *Elane^Cre^*, demonstrated that MCs arise from *Cpa3^+^* progenitors rather than *Ms4a3*^+^ or *Elane*^+^ GMPs. A corresponding population with a similar developmental trajectory was also identified in human early yolk sac and fetal liver, suggesting a conserved MC developmental program across species. These findings delineate the detailed developmental path of MCs in embryos, permitting future functional studies in immunity and development.

**Highlights:**

- *Cpa3*-expressing cells in the yolk sac and fetal liver contain mast cell precursors
- *Cpa3^CreERT2^* labels embryonic mast cells and their precursors
- Embryonic mast cells arise through *Cpa3^+^* mast cell precursors, but not *Elane*^+^/*Ms4a3^+^* GMPs
- The seeding of MCPs to embryonic skin slows down after E14.5

## Introduction

Mast cells (MCs) are immune sentinels that are widely distributed in connective and mucosal tissues. They participate in physiological and pathological processes, including type I allergic responses, tumor immune responses, and immunomodulation, by secreting a variety of bio-activators (Galli et al., 2020; St John et al., 2023). Therapeutic targeting of these cells may benefit the treatment of various diseases, such as atopic dermatitis, asthma and mastocytosis (Derakhshan et al., 2021; Valent et al., 2023), but requires deep understanding of their development.

Based on their tissue residency, murine MCs were classified into two main types, connective tissue MCs (CTMCs) and mucosal MCs (MMCs) (Derakhshan et al., 2021; Enerbäck, 1966; Katz et al., 1985). CTMCs locate mostly in the skin and peritoneal cavity, and their cytoplasmic granules contain heparin proteoglycan and relatively high concentrations of histamine, while MMCs locate mostly in gut and respiratory mucosa, and contain little or no heparin proteoglycans in their granules, and have lower amounts of histamine (Gurish and Austen, 2012; Katz et al., 1985). These two populations are with different origins and turnover rates (Chia et al., 2023; Tauber et al., 2023). Studies have shown that MCs could be generated both in embryos and adults (Gentek et al., 2018; Kitamura et al., 1979; Kitamura et al., 1977; Li et al., 2018), and MC precursors (MCPs) have been described in both embryos (Rodewald et al., 1996; Sonoda et al., 1983) and adults (Chen et al., 2005; Huang et al., 2016). Considerable work has been done to reveal MC development in adult bone marrow (BM), but its ontogeny is still under debate (Derakhshan et al., 2022). In adult BM, hematopoietic stem cells (HSCs) generate myeloid cells through multipotent progenitors (MPPs) and common myeloid progenitors (CMPs) (Akashi et al., 2000). CMPs further generate granulocyte-monocyte progenitors (GMPs), which were believed to give rise to all granulocytes and monocytes (Akashi et al., 2000). Studies showed that GMPs gave rise to MCs in *in vitro* culture (Arinobu et al., 2005). Subsequently, granulocyte progenitors (GPs), which are generated from GMPs, has been reported to generate neutrophils, eosinophils, basophils, and MCs (Sasaki et al., 2015). However, in other studies, MCPs with exclusive MC potential were identified as Lin^−^CD117^+^Sca1^−^Ly6C^−^FceR1^−^Integrin β7^+^T1/ST2^+^ in adult murine BM, which were derived from either MPPs or from CMPs but not from GMPs (Chen et al., 2005). Thus, fate mapping studies using fate mappers for GMPs and other progenitors are needed to reconcile these results. Although BM can undoubtedly produce MCs, its contribution seems to be limited to MCs in mucosal sites, such as gut, but not in CTMCs (Chia et al., 2023).

In contrast to MMCs, which could be replenished by BM-derived cells (Bankova et al., 2015; Derakhshan et al., 2021), CTMCs are rarely replenished by BM cells at steady state or in BM transplantation and parabiotic models (Dwyer et al., 2016; Franco et al., 2010; Gentek et al., 2018; Kitamura et al., 1979; Kitamura et al., 1977; Tauber et al., 2023). This evidence suggests that most CTMC populations, notably in the skin, are not generated by adult BM; rather, they arise from early embryonic hematopoiesis, seed tissues during organogenesis, and maintain themselves largely independently from adult BM (Gentek et al., 2018; Li et al., 2018). This pathway is similar to some tissue-resident macrophages, such as Langerhans cells (Hoeffel et al., 2012) and brain microglia (Ginhoux et al., 2010), as recently reviewed (St John et al., 2023). In agreement with this concept, a recent pan-organ single-cell transcriptomic analysis showed that *MrgprB2*^+^ CTMCs are embryonic hematopoiesis derived cells independent from the BM, while the *MrgprB2*^−^ MMCs develop after birth and are renewed by BM progenitors (Tauber et al., 2023). Collectively, MCs develop through two major paths, 1) most CTMCs are generated by embryonic hematopoiesis, and maintain themselves largely independently from adult BM, 2) MMCs are constantly generated by BM progenitors. Although considerable work has been done to reveal the development of MCs in adult BM, MC development in embryos is still incompletely understood, and it remains to be investigated whether embryonic MCs follow the same developmental path as BM-derived MCs.

During embryogenesis, three waves of hematopoiesis emerge sequentially in different sites (Hoeffel and Ginhoux, 2018). The earliest wave starts in the yolk sac (YS), an extra-embryonic membrane around the embryo (Hoeffel and Ginhoux, 2018). At embryonic day 7.0 (E7.0), the YS develops blood islands that initiate “primitive hematopoiesis” (Moore and Metcalf, 1970; Palis et al., 1999), generating primitive erythrocytes (nucleated erythrocytes), megakaryocytes, and macrophages (Palis et al., 1999; Xu et al., 2001). Later, the YS generates another wave of hematopoiesis, termed “transient definitive hematopoiesis”. The hematopoietic progenitor cells of this wave were initially termed erythro-myeloid progenitors (EMPs), as they produce mostly erythrocytes and myeloid cells (Frame et al., 2013; Gomez Perdiguero et al., 2015). Recent work has instead named them “late EMPs” to distinguish them from early cells with erythro-myeloid potential (“early EMPs”) generated during primitive hematopoiesis (Hoeffel et al., 2015). At E10.5, pre-HSCs produced in the aorta-gonads-mesonephros (AGM) region migrate to the fetal liver (FL) and begin “definitive hematopoiesis” (Boisset et al., 2010). Accordingly, Hoeffel *et al*. showed that, while early EMPs give rise to mostly microglia, late EMPs contribute major part of macrophages in tissues other than the central nervous system (Hoeffel et al., 2015). Similarly, MCs are also generated differentially by these hematopoiesis waves (Gentek et al., 2018; Li et al., 2018).

One study reported that MCs are first generated from YS primitive hematopoiesis and are then replaced by AGM-derived definitive hematopoiesis at a late stage of embryonic development (Gentek et al., 2018), while another study reported that MCs are primarily generated from late EMPs (transient definitive hematopoiesis) (Li et al., 2018). These embryonic MCs are abundant and functional, and could be primed by maternal IgE, leading to transient allergy in infants (Msallam et al., 2020). These studies revealed the embryonic origin and function of MCs, but the detailed developmental process and direct precursors for MCs remain largely unknown. A more comprehensive understanding of MC development in embryos would benefit therapeutic targeting of these cells, providing new options for treating allergies and other MC-associated immune dysfunctions.

Here, we used single-cell RNA sequencing (scRNA-seq), validated by fate mapping models, to analyze hematopoietic cells in the YS, FL, and embryonic skin. We identified a *Cpa3*-expressing population as early as E10.0 in the YS, and sequentially appeared in FL and embryonic skin from E10.5-E14.5, where they give rise to MCs within a temporal window that is mostly complete before E14.5 during the embryonic stage. Combined with in vitro differentiation assay and in vivo transplant method, we identified a Ter119^−^F4/80^−^CD45^+^CD117^+^CD16/32^+^CD135^−^CD115^−^Ly6C^−^ CD34^lo^ population in FL as potential MCPs. Furthermore, tracing with the GMP fate mappers *Ms4a3^Cre^* and *Elane^Cre^* revealed that MCs were derived not from FL GMPs but rather from a distinct lineage which can be marked by *Cpa3* expression. We also extended our findings to humans, identifying populations in the early human YS and FL that correspond to murine MCPs. These findings reveal a detailed developmental path of MCs in embryos and suggest a conserved MC developmental program across species.

## Results

### Single-cell RNA sequencing identifies *Cpa3*-expressing cells in E10.0 YS

Several MC populations, including skin MCs, were previously shown to be generated by embryonic hematopoiesis (Gentek et al., 2018; Li et al., 2018). Progenitors with MC potential have been identified in the YS at E9.5 using *in vivo* transplantation (Sonoda et al., 1983), but the MC committed progenitors and their trajectory to fetal MCs remained to be precisely characterized. To identify YS MCPs, we performed scRNA-seq on sorted GFP^+^ CD45^+^ and/or CD41^+^ hematopoietic cells from the YS of E10.0 GFP^+^ embryos. YS is in close contact with maternal tissues, thus easily to be contaminated by maternal cells. To exclude this contamination, we bred a GFP-transgenic (ubiquitously expressed) male (B6 ACTb-EGFP) with a C57BL/6 wild-type female, and used GFP^+^ embryos for cells sorting (**Figure 1A**). After filtering low-quality cells (see **Methods** for quality controls), we retrieved 5,990 cells for downstream analysis (**Figure 1A**). Unsupervised clustering identified twelve clusters that were annotated based on their differentially expressed genes (DEGs) and previously published datasets (Iturri et al., 2021; Tusi et al., 2018)(Ceccacci et al., 2023; Gao et al., 2022; Wang et al., 2020), including hematopoietic stem and progenitor cells (HSPCs) (c0_HSPC: *Hmga2*, *Cd34*, *Sox4*) (Tang et al., 2021; Yokomizo et al., 2022; Yokomizo et al., 2019), YS myeloid progenitors (YSMPs) (c1_YSMP: *Prtn3*, *Mpo*, *Hp*, *Plac8)*(Bian et al., 2020), macrophage precursors (c2_pMac) and macrophages (c3_Mac: *C1qa*, *C1qb*, *C1qc*, *Lpl*, *Dab2*, *Ccl9*) (Mass et al., 2016), megakaryocytes (Mk) (c9_Mk: *Myl9*, *Ppbp*, *Gp9*) and Mk progenitors (c6_pMk1: *Rap1b*, *Pbx1*, *F2r*) and precursors (c7_pMk2: *Timp3*, *Rab27b*, *Thbs1*, *Pf4*) (Iturri et al., 2021), erythrocytes (c8_Ery: *Hbb-bs*, *Hbb*-*a1*, *Hbb*-*a2*) and erythrocyte progenitors (c5_pEry: *Gm15915*, *Lmo2*, *Klf1*, *Car2*, *Blvrb*), endothelial cells (c10_Endo: *Lyve1*, *Mest*, *Sparc*, *Plvap*), and fibroblasts (c11_Fibroblast: *Col1a1*, *Col1a2*, *Col3a1*) (Iturri et al., 2021; Tusi et al., 2018) (**Figure 1B**). Notably, one cell cluster can be defined by its prominent expression of *Cpa3, Perp* and *Gata2* (**Figures 1B-D**), three genes known to be expressed by MCs and basophils (**Figures S1A and S1B**) (Jippo et al., 1996; Tauber et al., 2023). These data suggested that this *Cpa3*-expressing cluster might contain potential basophil precursors (BaPs) and MCPs in the YS.

**Figure 1.**
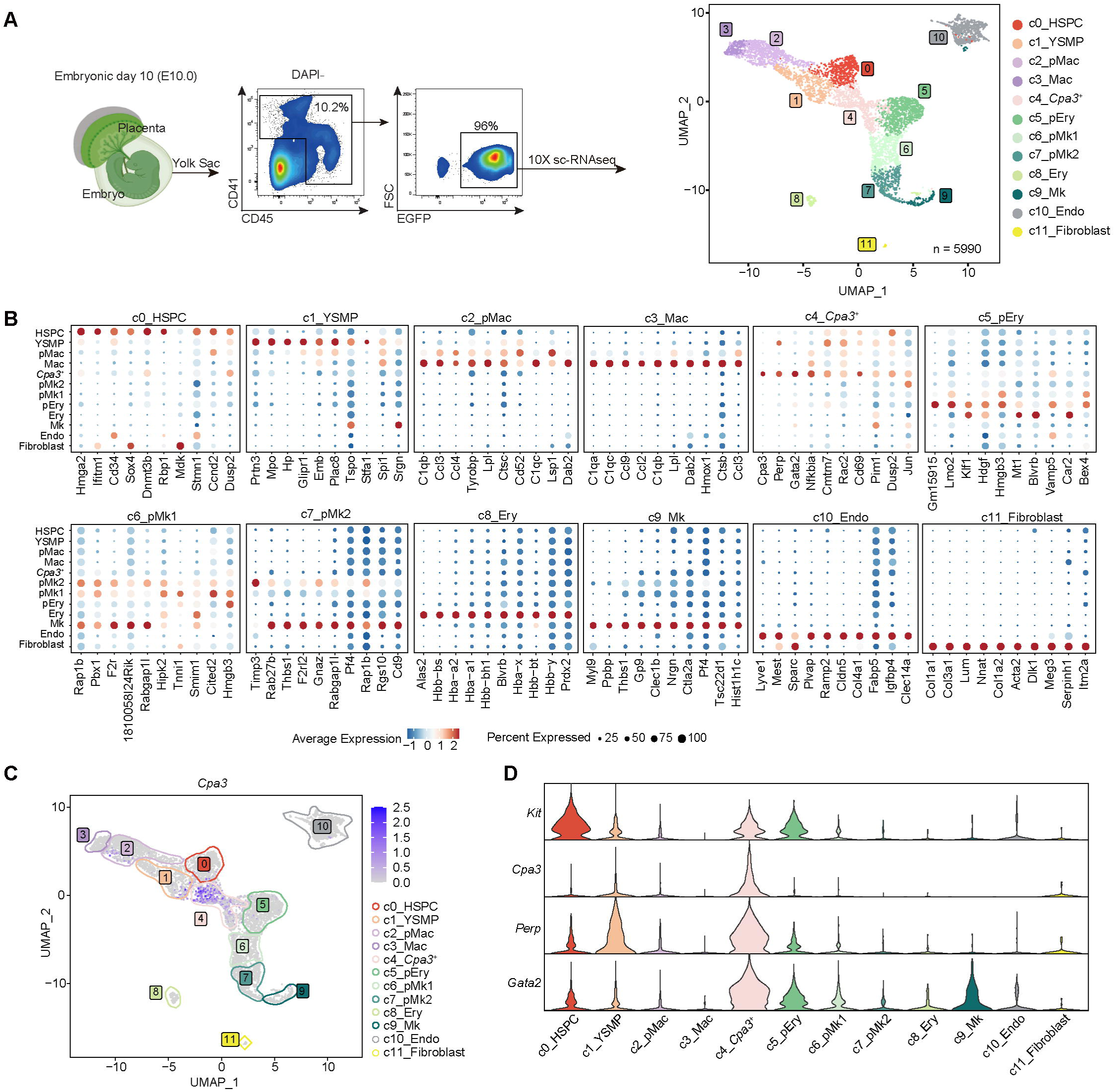
Single-cell RNA sequencing identifies *Cpa3*-expressing cells in E10.0 YS. (A) Schematic (left) showing the workflow of scRNA-seq. CD41^+^ and/or CD45^+^ EGFP^+^ cells were sorted from the YS of E10.0 *EGFP^+^* embryos, and single-cell transcriptomes were generated using 10x Genomics workflow. Uniform Manifold Approximation and Projection (UMAP) plot (right) of the scRNA-seq performed on sorted cells from the YS (n = 5,990). (B) Dot plot showing the expression of top 10 DEGs across clusters identified in (A). Colors indicate the average expression of each gene. Spot sizes represent the proportion of gene-expressing cells. (C) Feature plot showing the expression of *Cpa3* among the clusters identified in (A). (D) Violin plots showing the expression of selected genes among the top markers for *Cpa3*-expressing cells and the known MC signature gene *Kit*.

### Fate mapping with *Cpa3^CreERT2^* model reveals that *Cpa3*^+^ progenitors give rise to MCs

To understand the fate of these *Cpa3*^+^ cells *in vivo*, we generated a tamoxifen-inducible fate mapping mouse model using a *Cpa3^CreERT2^* mouse line (**Figure S2A**) crossed to a *Rosa^tdTomato^* line (Madisen et al., 2010). First, to validate the existence of *Cpa3*-expressing cells in the YS, C57BL/6 WT females intercrossed with *Cpa3^CreERT2^-Rosa^tdTomato^* males were administered a single dose of tamoxifen at E9.5 and analyzed at E10.5. Validating our above-mentioned observation of a *Cpa3*^+^ cluster in the YS, tdTomato^+^ cells, although very few in numbers, could be detected in the YS, but not in the embryo proper (**Figure S2B**). We then traced these *Cpa3*-expressing YS cells by administering a single dose of tamoxifen at E10.5 and analyzed tdTomato^+^ cells in the *Cpa3^CreERT2^-Rosa^tdTomato^* embryos at different timepoints. At E11.5, most of the tdTomato^+^ cells were in the FL, with a small number in the body and a negligible number in the YS (**Figure 2A**). Significantly more tdTomato^+^ cells were observed in the FL and skin at E12.5 whereas YS exhibited very few tdTomato^+^ cells indicating decreased output of Cpa3^+^ cells from YS at this stage (**Figure 2B**). At E14.5 and E16.5, tdTomato^+^ cells were mostly found in the skin and FL, with a small number detected in the lung, kidney, and gut (**Figures 2C and S2C**). Thus, we focused on FL and skin hereafter.

**Figure 2.**
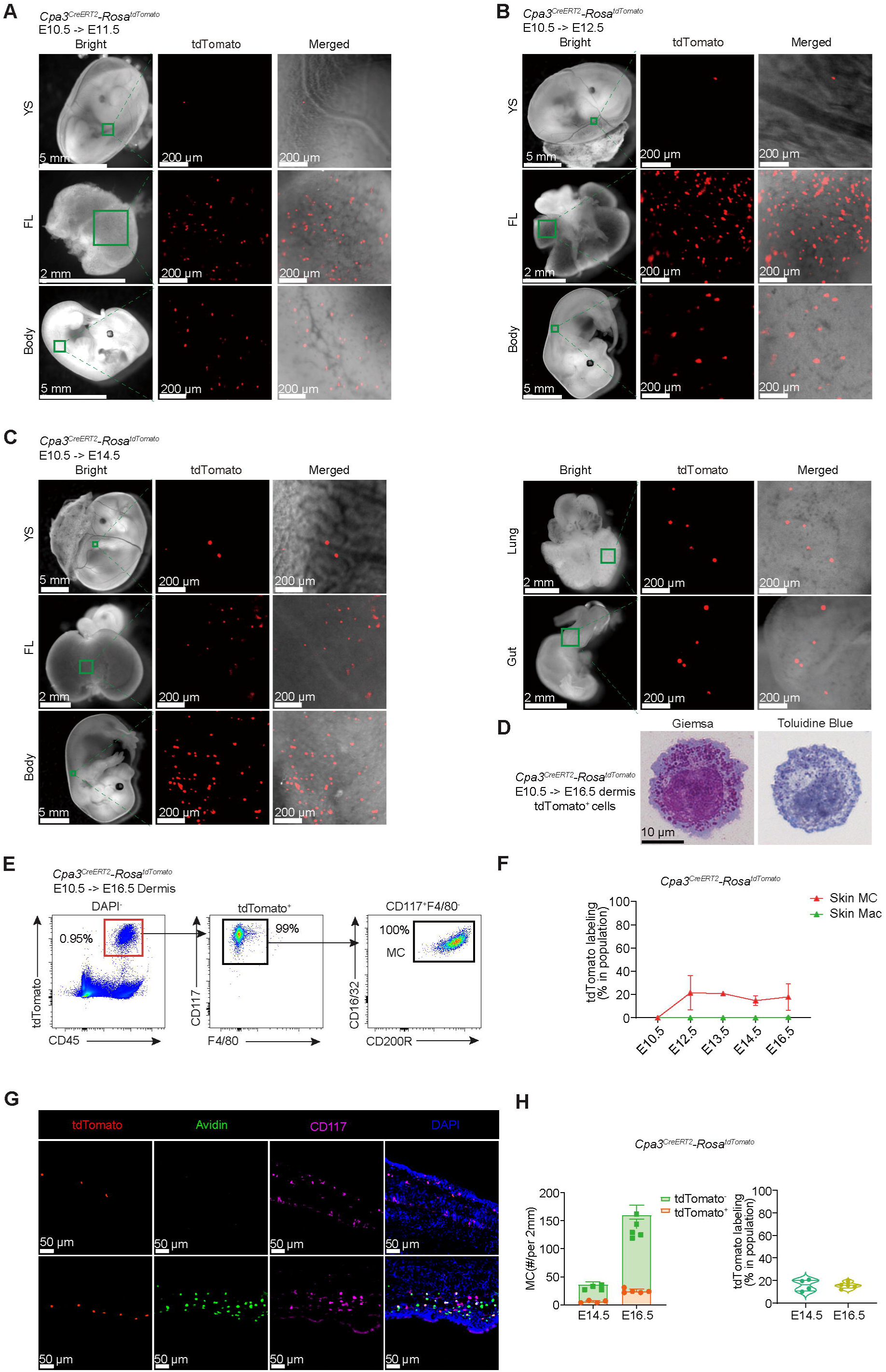
Fate mapping with *Cpa3^CreERT2^*model reveals that *Cpa3*^+^ progenitors give rise to MCs. (A-C) Representative brightfield and fluorescence microscopy images showing E11.5 (A), E12.5 (B), and E14.5 (C) *Cpa3^CreERT2^-Rosa^tdTomato^*embryos pulsed with tamoxifen at E10.5. Red fluorescence indicates tdTomato^+^ cells. (D) Representative brightfield microscopy images showing the morphology (stained with Giemsa or toluidine blue) of fate mapped tdTomato^+^ cells that sorted from the skin of E16.5 *Cpa3^CreERT2^-Rosa^tdTomato^*embryos pulsed with tamoxifen at E10.5. (E) Flow cytometry analysis of tdTomato^+^CD45^+^ cells in the dermis of E16.5 *Cpa3^CreERT2^-Rosa^tdTomato^* embryos pulsed with tamoxifen at E10.5. (F) Percentages of tdTomato-expressing MCs in dermis after tamoxifen induction at E10.5. (G, H) Confocal immunofluorescence microscopy images showing tdTomato^+^ cells in the skin of E14.5 and E16.5 *Cpa3^CreERT2^-Rosa^tdTomato^* embryos pulsed with tamoxifen at E10.5. Scale bar = 50 μm (G). Numbers of tdTomato^+^ and tdTomato^−^ MCs in 2 mm back skin of embryo (H left, and Figure S2D) and the percentages of tdTomato-expressing MCs over total MCs (H right) of E14.5 (n = 4) and E16.5 (n = 5) *Cpa3^CreERT2^-Rosa^tdTomato^* embryos pulsed with tamoxifen at E10.5.

Next, we performed Giemsa and toluidine blue staining to clarify the identity of these tdTomato^+^ cells in E16.5 skin. We found that these cells contained many granules (**Figure 2D**), morphologically resemble MCs. In addition, almost all the tdTomato^+^ cells in E16.5 dermis were CD117^+^CD16/32^+^, an MC phenotype (Gentek et al., 2018) (**Figure 2E**). These tdTomato^+^ cells comprised up to 20% of total CD117^+^CD16/32^+^ MCs in the embryonic skin (**Figure 2F**). Immunofluorescence imaging also showed that these tdTomato^+^ cells were CD117^+^ at E14.5 and CD117^+^Avidin^+^ at E16.5 (**Figures 2G, 2H and S2D**), confirming their MC identity and their maturation along developmental timepoints, as indicated by the accumulation of heparin-containing granules that can be stained with fluorophore-conjugated avidin. To validate if these tdTomato^+^ cells were able to give rise to MCs, we sorted tdTomato^+^CD45^+^ cells from E13.5 FL of *Cpa3^CreERT2^-Rosa^tdTomato^*embryos that were pulsed with tamoxifen at E11.5, and transferred into peritoneal cavity of sash (*Kit^W-sh/W-sh^*) mice (Grimbaldeston et al., 2005), which lack MCs in all tissues thus have open “niches” for transferred cells to differentiate into MCs, and analyzed 2 weeks after transfer. Donor-derived tdTomato*^+^* MCs could be detected in the peritoneal lavage of the recipients (**Figure S2E)**. Collectively, fate mapping with the *Cpa3^CreERT2^* model demonstrated that *Cpa3*-expressing FL cells contain precursors that give rise to skin MCs.

### Single-cell RNA sequencing identifies MCPs in the FL

In *Cpa3^CreERT2^-Rosa^tdTomato^* embryos pulsed with tamoxifen at E10.5, tdTomato^+^ cells mostly appeared in the FL at E11.5 (**Figure 2A**). To fully investigate the nature of these cells, we performed scRNA-seq for tdTomato^+^ (n = 7,232) and tdTomato^−^CD45^+^ (n = 7,701) cells from the FL of E13.5 *Cpa3^CreERT2^-Rosa^tdTomato^*mice pulsed with tamoxifen at E11.5 (**Figure 3A**). Unsupervised clustering identified 11 clusters that were annotated based on their DEGs and previously published datasets (Ceccacci et al., 2023; Gao et al., 2022; Wang et al., 2020), including early progenitors (c0_Progenitor: *Cd34*, *Hmga2*, *Sox4*, *Ctla2a*) (Sarah et al., 2020), MEPs (c2_MEP: *Vamp5*, *Ass1*, *Car2*), erythrocytes (c1_Ery: *Hba-a1*, *Hba-a2*, *Cpox*, *Blvrb*, *Alad*) and megakaryocytes (c9_Mk: *Pf4*, *Rab27b*, *Plek*, *Rap1b*), neutrophils (c8_Neu: *Retnlg*, *Stfa2*, *Ltf*, *S100a9*) and neutrophil precursors (c3_NeuP: *Prtn3*, *Ctsg*, *S100a9*, *S100a8*), monocyte and dendritic cell (DC) precursors (MDPs) (c5_MDP: *Ifitm3*, *Ctsc*, *Psap*, *Irf8*), macrophages (c6_Mac: *C1qc*, *C1qa*, *C1qb*, *Marco*, *Lgmn*, *Ctsb*, *Hmox1*), and eosinophils (c10_Eo: *Prg3*, *Ear1*, *Ear6*) (**Figure 3B**) (Ceccacci et al., 2023; Gao et al., 2022; Wang et al., 2020). As in the YS, the FL contained a *Cpa3*-expressing cluster (c4_*Cpa3*^+^) that also expressed *Gata2* (**Figure 3B**).

**Figure 3.**
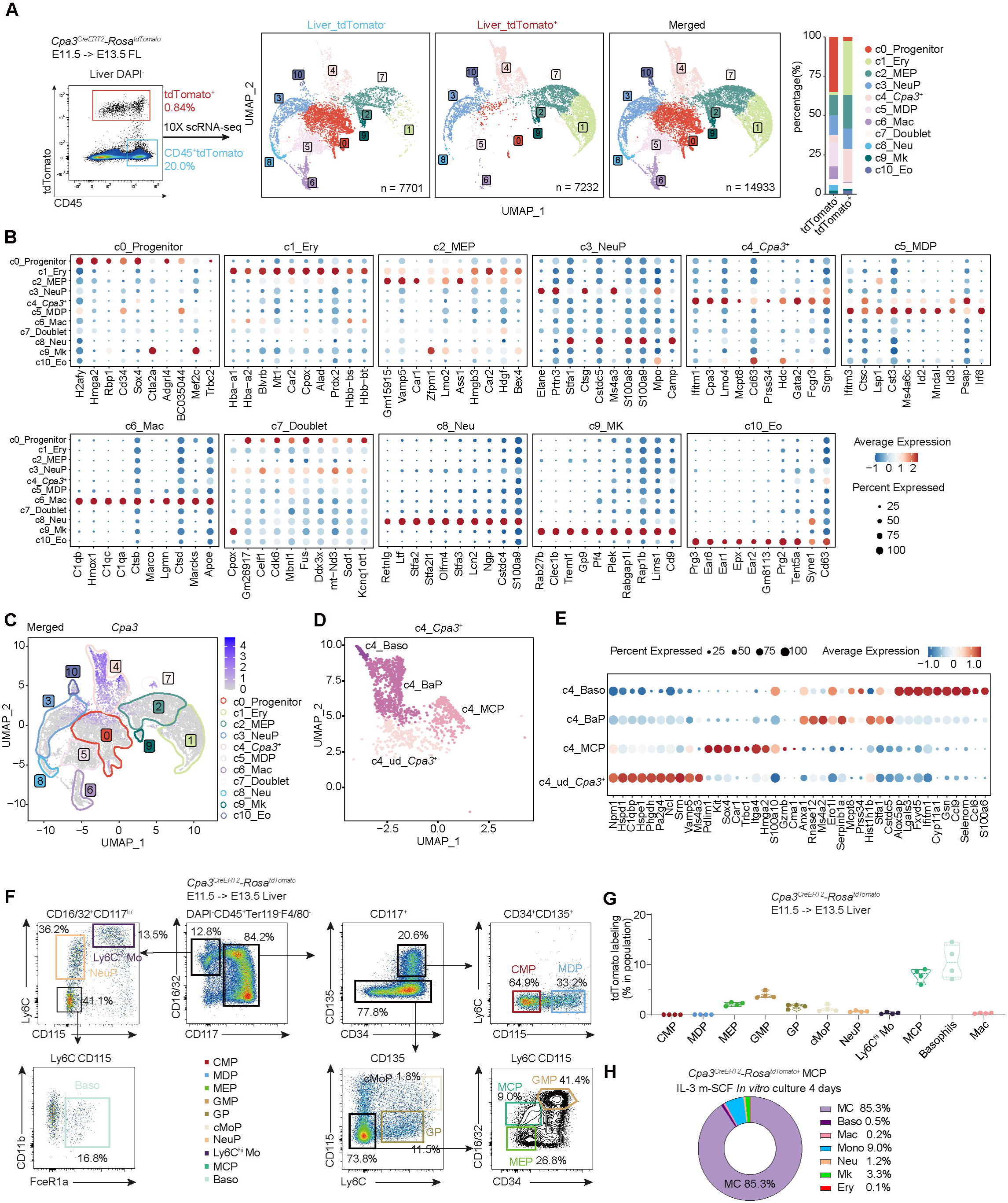
Single-cell RNA sequencing identifies MCPs in the FL. (A) Schematic (left) showing the workflow of scRNA-seq for tdTomato^+^ (n = 7,232) and tdTomato^−^CD45^+^ (n = 7,701) cells from the FL of E13.5 *Cpa3^CreERT2^-Rosa^tdTomato^* embryos pulsed with tamoxifen at E11.5. UMAP plots showing population in merged and separated cells fractions. Adjacent bar charts showing the proportions of each cluster within the tdTomato^+^ and tdTomato^−^ CD45^+^ fractions. (B) Dot plots showing the top 10 DEGs across clusters identified in the FL. Colors indicate the average expression of each gene. Spot sizes represent the proportion of gene-expressing cells. (C) Feature plot showing the expression of *Cpa3* in FL scRNA-seq data. (D, E) UMAP plot (D) showing the heterogeneous subclusters identified within the *Cpa3*^+^ cluster. Dot plot (E) showing the expression of top 10 DEGs for each subcluster. (F, G) Representative flow cytometry plots (F) showing the gating strategy for FL GMPs and MCPs in E13.5 *Cpa3^CreERT2^-Rosa^tdTomato^* embryos pulsed with tamoxifen at E11.5. Histogram (G) showing the labeling across the populations identified in (F). (H) The percentage of progenies of *Cpa3^CreERT2^-Rosa^tdTomato+^*MCPs after 4 days cultured in medium contained IL-3 and murine stem cell factor (m-SCF).

The tdTomato^+^ fraction contained mostly c1_Ery, c2_MEP, c4_*Cpa3^+^*, and a fraction of c10_Eo and c3_NeuP (**Figure 3A**), likely due to some expression of *Cpa3* in c0_Progenitor (**Figure 3C**). Of note, not all the tdTomato^+^ cells express *Cpa3*, MEPs, erythrocytes and Mks showed negligible expression of *Cpa3*, suggesting that the tdTomato labeling of these cells occurred earlier at their *Cpa3^+^* precursors (**Figure S3A**). The c0_Progenitor (**Figures S3B and S3C**) and c4_*Cpa3^+^*(**Figures 3D and 3E**) clusters appeared to be heterogeneous and were thus further subclustered (**Figure S3D**). The early progenitors could be further divided into four subclusters: two HSPC subclusters (*Hlf*), with one subcluster expresses cell cycle gene (*Top2a*, *Ube2c*), and common lymphoid progenitor (CLP) (*Il7r*, *Dntt*) and GMP (*Mpo*, *Elane*) subclusters (**Figures S3B and S3C**). The c4_*Cpa3^+^* cluster could also be divided into four subclusters: basophil precursor (BaP) (*Serpinb1a*, *Ms4a2*), basophil (*Mcpt8*, *Prss34*, *Lgals3*, *Alox5ap*), MCP (*Car1*, *Sox4*, *Kit*, *Itga4*) (Dahlin et al., 2018), and a cluster located close to c0_progenitor (**Figure S3D**) that might represent bipotential or undifferentiated progenitors for basophils and MCs, thus were termed undifferentiated *Cpa3*^+^ progenitors (ud_*Cpa3*^+^) (**Figures 3D and 3E**).

In adult BM, MCPs were identified as Lin (CD3, CD4, CD8, CD11b, B220, Gr-1 and Ter119)^−^ CD117^+^Sca1^−^Ly6C^−^FceR1^−^Integrin β7^+^T1/ST2^+^ (Chen et al., 2005). In FL, MCPs were identified as Integrin β7^+^CD117^+^CD11b^low^ precursors (Li et al., 2018). Profiling the expression of those genes in FL scRNA-seq data showed that *Itgb7* (encodes Integrin β7) and *Il1rl1* (encodes ST2) were expressed by only a small fraction of MCPs, thus they were not able to be used to define the whole population (**Figure S3E**). E-cadherin was reported to mark basophil and MC precursors in adult BM (Wanet et al., 2021). In FL, *Cdh1* (encodes E-cadherin) was expressed by some BaPs but not MCPs (**Figure S3F**), suggesting a difference between embryonic and adult hematopoiesis. Thus, a new gating strategy is needed to better characterize MCPs in FL.

To develop a new gating strategy for flow cytometric analysis of FL cells, we referred to several gating strategies described in previous studies (Liu et al., 2019; Yanez et al., 2017) and the surface marker gene expression profile from scRNA-seq data, to map the populations described above (**Figure S3G**), and we found that GMPs expressed a higher level of *Cd34* than MCPs (**Figures S3G and S3H**). For FL DAPI^−^Ter119^−^CD45^+^ cells, macrophages were identified as the F4/80^+^ fraction, while F4/80^−^ fraction contained progenitors which were defined as CD117^+^ and differentiated myeloid cells which were defined as CD117^−^CD16/32^+^. In the progenitor fraction (CD117^+^), CD135^+^CD34^+^ cells contained mostly CMPs (CD117^+^CD34^+^CD135^+^CD115^−^Ly6C^−^) and MDPs (CD117^+^CD34^+^CD135^+^CD115^+^Ly6C^−^), while CD135^−^ cells contained common monocyte progenitors (cMoPs) (CD117^+^CD16/32^+^Ly6C^hi^CD115^+^), granulocyte progenitors (GPs) (CD117^+^CD16/32^+^Ly6C^+^CD115^−^), and some Ly6C^−^ cells including GMPs (CD117^+^CD16/32^+^CD34^+^CD135^−^Ly6C^−^CD11b^−^), MEPs (CD117^+^CD16/32^−^CD34^−^CD135^−^), and MCPs (CD117^+^CD16/32^+^CD34^lo^CD135^−^Ly6C^−^CD11b^−^) (Chen et al., 2005; Hoeffel et al., 2015; Liu et al., 2019; Wanet et al., 2021; Yanez et al., 2017; Yanez et al., 2015). Finally, the CD117^−^ CD16/32^+^ differentiated myeloid cells contained monocytes (Lin^−^CD117^−^ CD16/32^+^Ly6C^hi^CD115^+^), NeuPs (Lin^−^CD117^−^CD16/32^+^Ly6C^int^CD115^−^), and basophils (Lin^−^ CD117^−^CD16/32^+^Ly6C^−^CD115^−^CD11b^−^FceR1^+^) (Matsumura et al., 2022; Wanet et al., 2021) (**Figure 3F**).

Using the gating strategy above (**Figure 3F**), we analyzed tdTomato^+^ labeling of cell populations in the FL of E13.5 *Cpa3^CreERT2^-Rosa^tdTomato^*mice pulsed with one dose of tamoxifen at E11.5. MCPs and basophils showed the highest labeling across these populations (**Figures 3F and 3G**). Analysis by gating on tdTomato*^+^* cells showed that they contained both CD45^+^ and CD45^−^CD117^+^ (erythrocyte progenitors); the CD45^+^ fraction contained MEPs, GMPs, MCPs, a few basophils, and neutrophils (**Figure S3I**). This minor labeling of neutrophils and monocytes might be due to some non-specific expression of *Cpa3* in GMPs, as seen in the scRNA-seq data (**Figure 3C**). To further validate that these tdTomato^+^ cells in our MCP gate were with indicated potential, we sorted tdTomato*^+^* MCPs from the FL of E13.5 *Cpa3^CreERT2^-Rosa^tdTomato^* embryos pulsed with tamoxifen at E11.5, for *in vitro* differentiation for 4 days (**Figure 3H**), demonstrating that these cells gave rise to MCs. Collectively, these data demonstrate that FL MCPs could be defined as Ter119^−^F4/80^−^CD45^+^CD117^+^CD16/32^+^CD135^−^CD115^−^Ly6C^−^CD34^lo^.

### Embryonic MCs do not arise from *Ms4a3*^+^ or *Elane*^+^ GMPs

We next performed pseudotime trajectory analysis of the above FL scRNA-seq data to predict the developmental trajectory of MCs using the Monocle 3 toolkit (Cao et al., 2019). In FL, the HSPC-rooted trajectory separated to three branches developing into basophils/MCPs/MEPs/erythrocytes/megakaryocytes, GMPs/neutrophils/monocytes/macrophages, and lymphoid cells respectively (**Figure 4A**), in agreement with previous reports in both human and mice (Popescu et al., 2019; Tusi et al., 2018; Wang et al., 2020). Trajectory analyses in FL suggested that MC lineage separated from neutrophil and monocyte lineages at an early stage, indicating that MCPs might not arise from GMPs.

**Figure 4.**
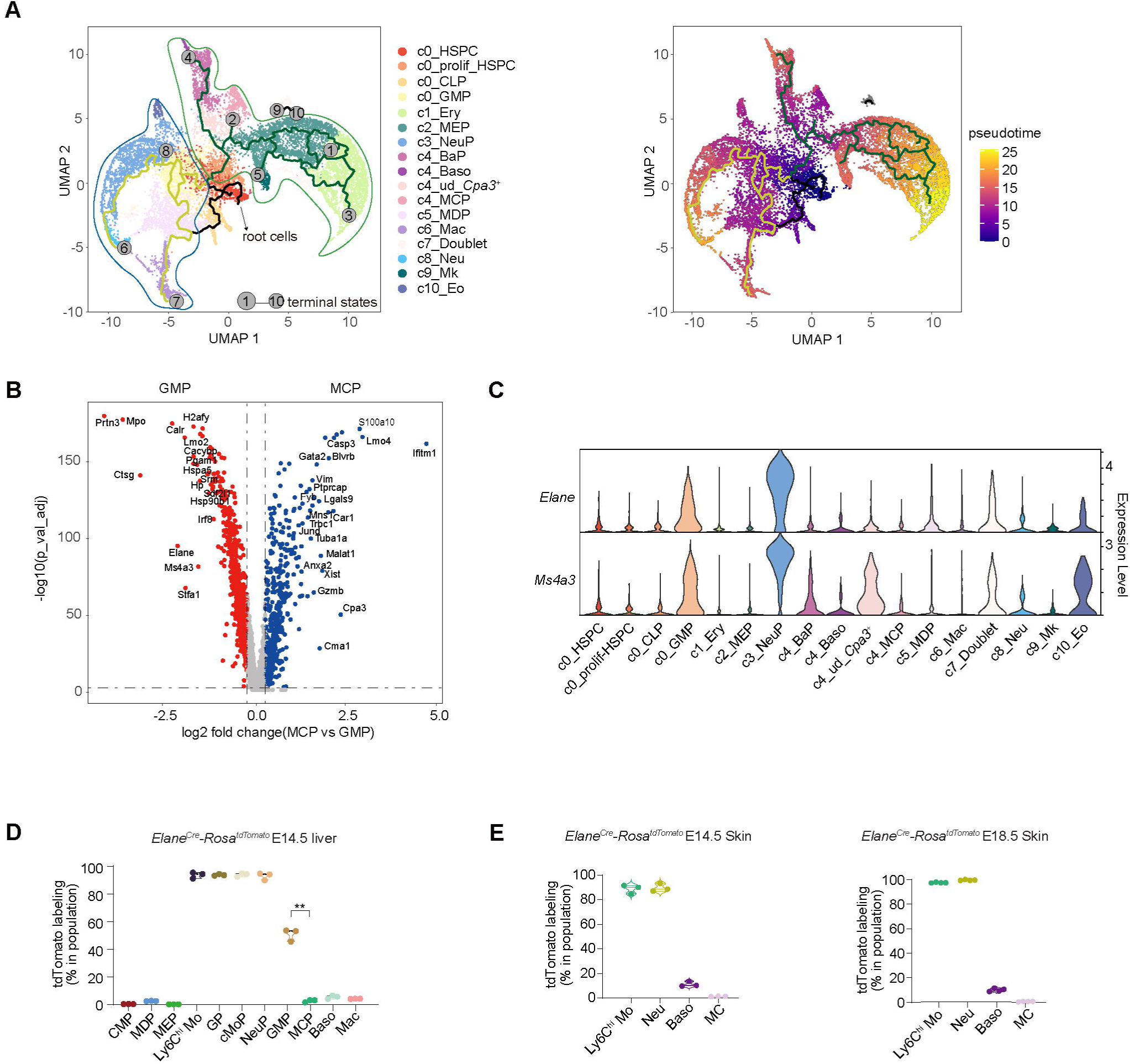
MCs do not arise from *Ms4a3*^+^ or *Elane*^+^ GMPs. (A) Inferred pseudotime trajectories of cell development in the FL by monocle3. Lines indicate inferred developmental trajectories. Cells are colored according to identity clusters (left). UMAP plots (right) showing cells colored by pseudotime. (B) Volcano plot showing the DEGs between GMPs and MCPs. Genes in red are upregulated in GMPs; genes in blue are upregulated in MCPs. Vertical dashed lines indicated Log2 fold changes = −0.25 or 0.25, horizontal dashed line = −log_10_(0.05). (C) Violin plots showing the expression of *Ms4a3* and *Elane* within the clusters identified in the FL scRNA-seq dataset shown in Figure S3D. (D) Percentage of tdTomato-expressing cells within the indicated populations in the FL of E14.5 *Elane^Cre^-Rosa^tdTomato^* embryos (n = 3). (E) Percentage of tdTomato-expressing cells within the indicated populations in the dermis and brain of E14.5 (n = 3) and E18.5 (n = 4) *Elane^Cre^-Rosa^tdTomato^* embryos.

To validate this predicted separation of MCP and GMP lineages, we began by analyzing the DEGs between GMPs vs. MCPs (**Figure 4B**). *Ms4a3* and *Elane* appeared in the top 20 DEGs that were highly expressed in GMPs but not MCPs (**Figures 4B and S4A, S4B**). We previously generated an *Ms4a3^Cre^-Rosa^tdTomato^* GMP fate mapping mouse model which labels GMPs and their progeny including granulocytes and monocytes (Liu et al., 2019). In adult *Ms4a3^Cre^-Rosa^tdTomato^* mice, neutrophils and basophils were highly labeled (>90%), while MCs were not (<10%), suggesting that MCs were not the progeny of GMPs, consistent with a recent report (Tauber et al., 2023). Furthermore, in the FL and dermis of E14.5 *Ms4a3^Cre^-Rosa^tdTomato^* embryos, while FL GMPs and their progenies were labeled, liver MCPs and dermal MCs were not (**Figures S4C and S4D**), demonstrating that MCs were not derived from *Ms4a3*^+^ GMPs. Notably, although MCs were not labeled, 22% of basophils were labeled (**Figure S4D**), likely due to the expression of *Ms4a3* in BaPs (**Figure S4A**). *Elane* was another DEG that highly expressed by GMPs but not by MCPs. Differ from *Ms4a3*, *Elane* was not expressed by BaPs (**Figures 4C and S4B**). We thus used an *Elane^Cre^-Rosa^tdTomato^* model (**Figure S4E**) to further separate MCP and GMP lineages. In E14.5 *Elane^Cre^-Rosa^tdTomato^*embryos, GMPs, and NeuPs in the FL were highly labeled, while MCPs in the FL and MCs and basophils in the dermis were not (**Figures 4D and 4E**), confirming an early separation between the MCP and GMP lineages. Collectively, combining computational trajectory analysis with *Ms4a3^Cre^* and *Elane^Cre^* GMP fate mapping models, we demonstrated that MCs are not the progeny of GMPs, but instead they are derived from the *Cpa3*-expressing MCPs of an independent lineage.

### The seeding of MCPs to embryonic skin slows down after E14.5

To fully investigate the cells mapped in the peripheral tissues of *Cpa3^CreERT2^-Rosa^tdTomato^* embryos, we performed scRNA-seq for tdTomato^+^ (n = 4,314) and tdTomato^−^CD45^+^ (n = 11,763) cells from the body wall of E13.5 *Cpa3^CreERT2^-Rosa^tdTomato^* embryos pulsed with tamoxifen at E11.5 (**Figure 5A**). Unsupervised clustering identified 11 clusters that were manually annotated based on their DEGs, including two macrophage populations (c0_Mac_1: *C1qa*, *C1qb*, *C1qc*, *Mrc1*, *Dab2*, *Apoe*; c8_Mac_2: *Sparc, Serpine2, Olfml3, P2ry12*), and a proliferating macrophage population (c6_prolif_Mac: *Top2a*, *Mki67*); monocytes and DCs (c1_Mo&DC: *S100a6*, *Cd52*); lymphoid cells (c4_Lymphoid: *Il7r*); and neutrophils (c5_Neu: *S100a9*, *S100a8*), endothelia cells (c9_Endo: *Nnat, Meg3, Ptn, Sox11,Stmn2*) and erythrocytes (c10_Ery: *Hbb-bh1, Hbb-a1, Hbb-bt, Hbb-a2, Hbb-bs*) (**Figure 5B**). As in the YS and FL, *Cpa3*^+^ cells could also be observed in the dermis, containing mostly MCs (c2_MC: *Srgn*, *Gata2*, *Hdc*, *Cma1*, *Tpsb2*), proliferating MCs (c3_prolif_MC: *Casp3*, *Tpsg1*), and basophils (c7_Baso: *Ccl9, Mcpt8*, *Cd200r3*, and *Mpc2*) (**Figures 5A, 5B, S5A**) (Cohen et al., 2018; Jacob et al., 2023; Tauber et al., 2023). Importantly, the tdTomato^+^ fraction was composed mainly of MCs and basophils (**Figure 5A**).

**Figure 5.**
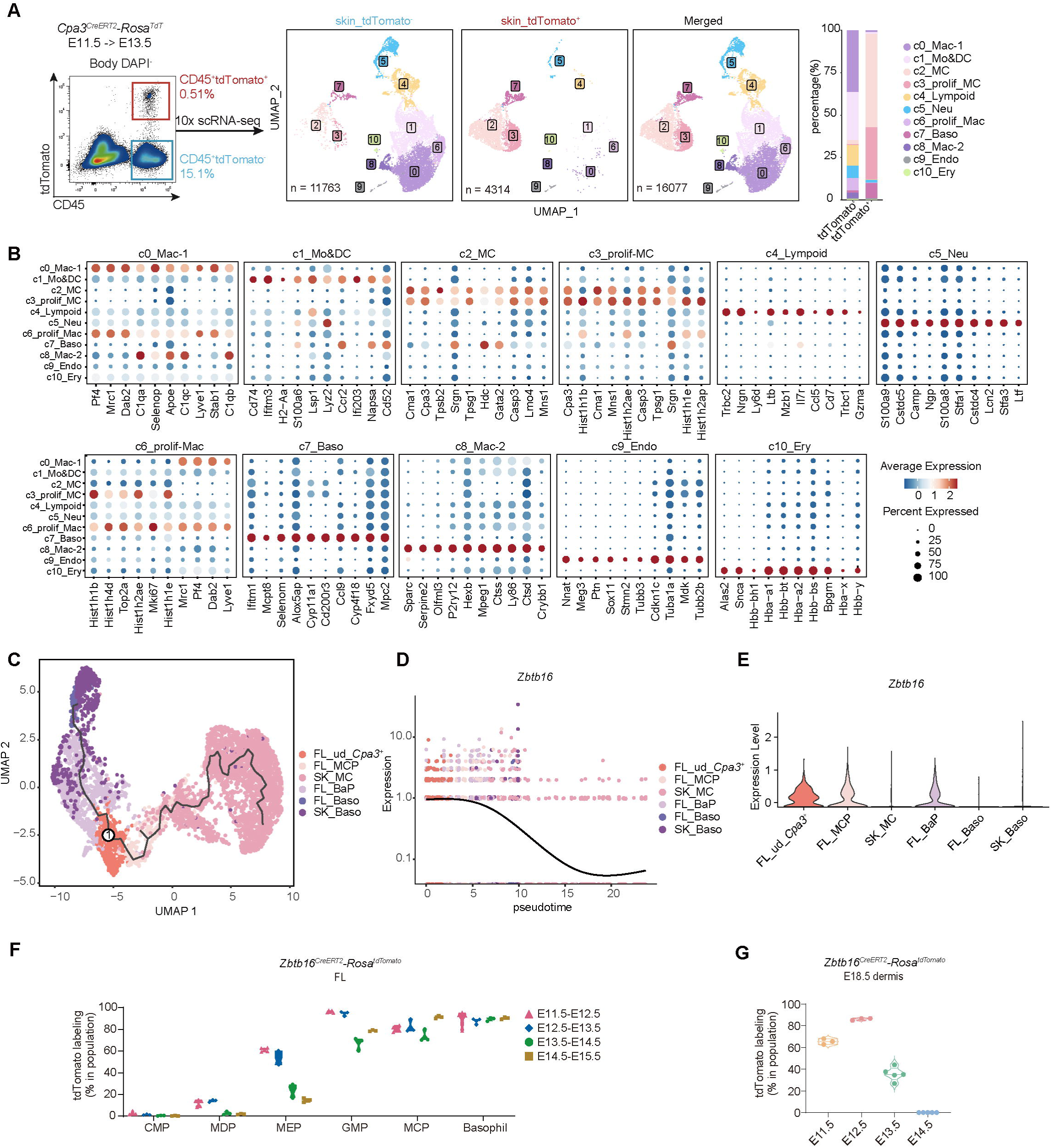
The seeding of MCPs to embryonic skin slows down after E14.5. (A) Schematic showing the workflow of scRNA-seq for tdTomato^+^ (n = 4,314) and tdTomato^−^ CD45^+^ (n = 11,763) cells from the bodies of E13.5 *Cpa3^CreERT2^-Rosa^tdTomato^*embryos pulsed with tamoxifen at E11.5. UMAP plots showing population in merged and separated cells fractions. Adjacent bar charts showing the proportions of each cluster within tdTomato^−^CD45^+^ the and tdTomato^+^ fractions. (B) Dot plots showing the top 10 DEGs across the clusters identified in (A). Colors indicate the average expression of each gene. Spot sizes represent the proportion of gene-expressing cells. (C) UMAP plot showing Inferred pseudotime trajectories of the merged *Cpa3*^+^ cells in the FL and dermis. Lines indicate inferred developmental trajectories. (D) Expression of *Zbtb16* over pseudotime, inferred by monocle3. (E) Violin plot showing the expression of *Zbtb16*. (F) Percentage of tdTomato-expressing cells in each population in the FL of E12.5 (yellow, n = 5), E13.5 (red, n = 3), E14.5 (green, n = 3) and E15.5 (blue, n = 3) *Zbtb16^CreERT2^-Rosa^tdT^* embryos pulsed with tamoxifen one day before each timepoint. (G) Percentage of tdTomato-expressing MCs and basophils in the dermis of E18.5 *Zbtb16^CreERT2^-Rosa^tdT^* embryos pulsed with tamoxifen at E11.5 (n = 3), E12.5 (n = 3), E13.5 (n =5), or E14.5 (n = 5).

To uncover the path from FL *Cpa3*^+^ precursors to those *Cpa3*^+^ MCs in the dermis, we merged scRNA-seq datasets from the FL (c4_*Cpa3^+^* cluster) and the dermis (c2_MC and c7_Baso clusters) and performed trajectory analysis (**Figure 5C**). This revealed that basophils and MCs emerged as two separate branches from ud_*Cpa3*^+^ (**Figure 5C**). To address the time window for dermal MC generation, we aimed to find a gene that was expressed transiently in MCPs. To do so, we profiled the expression of DEGs between FL MCPs and skin MCs (**Figure S5B**) over pseudotime and identified *Zbtb16* as a highly expressed gene by MCPs but not mature MCs (**Figures 5D, 5E, and S5C**), suggesting that *Zbtb16* was expressed transiently by MCPs but downregulated in terminally differentiated basophils and MCs. Using a *Zbtb16^CreERT2^-Rosa^tdTomato^* tamoxifen-inducible fate mapping model (**Figure S5D**) to label MCPs, it enabled us to investigate their development into mature MCs. Tamoxifen induction at E11.5 efficiently labeled FL MCPs (80.8 ± 3.47%) at E12.5 (**Figure 5F**) and subsequently dermal MCs (60.9 ± 12.6%) at E18.5 (**Figure 5G**). However, tamoxifen induction at E14.5 labeled FL MCPs at E15.5 but not dermal MCs at E18.5 (**Figure 5G**), suggesting that MCPs generated after E14.5 did not contribute significantly to dermal MCs. Thus, E18.5 dermal MCs largely arose from *Zbtb16*^+^ progenitors generated before E14.5. Collectively, our analyses revealed dynamic changes alongside the development of FL MCPs into MCs in peripheral tissues and demonstrated that MCPs seed dermis in an embryonic development window, largely from E11.5 (the timepoint at which MC progenitors can be detected in the *Cpa3^CreERT2^-Rosa^tdTomato^*embryo proper) to E14.5 (the timepoint at which giving tamoxifen to *Zbtb16^CreERT2^-Rosa^tdTomato^* embryos does not label dermal MCs), and slow down thereafter.

### MC development is conserved across species

To address whether MC development is conserved between humans and mice, we explored the published human YS (Goh et al., 2023) and FL (Popescu et al., 2019) scRNA-seq datasets. The human YS also contains *CPA3*-expressing cells that were annotated as human MCs or eosinophil-basophils (**Figures 6A and S6A**). Human MEMPs also showed a low level of *CPA3* expression (**Figure 6A**). The population of annotated MCs did not express the mature MC marker *CMA1* (**Figure S6B**), suggesting that they might at precursor stage of MCs. These populations showed great similarity to murine YS *Cpa3*^+^ cells in Pearson correlation analysis (**Figure 6B**). Next, we explored the corresponding *CPA3*-expressing cell population in human FL. Here, *CPA3* expression was also detected in MCs and MEMPs (**Figures 6C and S6C**). Similar with their YS counterparts, FL MCs did not express *CMA1* (**Figure S6D**), suggesting that these cells might at precursor stage of MCs. In correlation analysis, these two populations showed great similarity to murine MCPs (**Figure 6D**). Finally, trajectory analysis showed that these human MCPs separated from other granulocyte lineages at the stage before GMP in both YS (**Figure 6E**) and FL (**Figure 6F**), confirming the result of Popescu et al (Popescu et al., 2019). Together, these analyses suggest that MC development is conserved across species.

**Figure 6.**
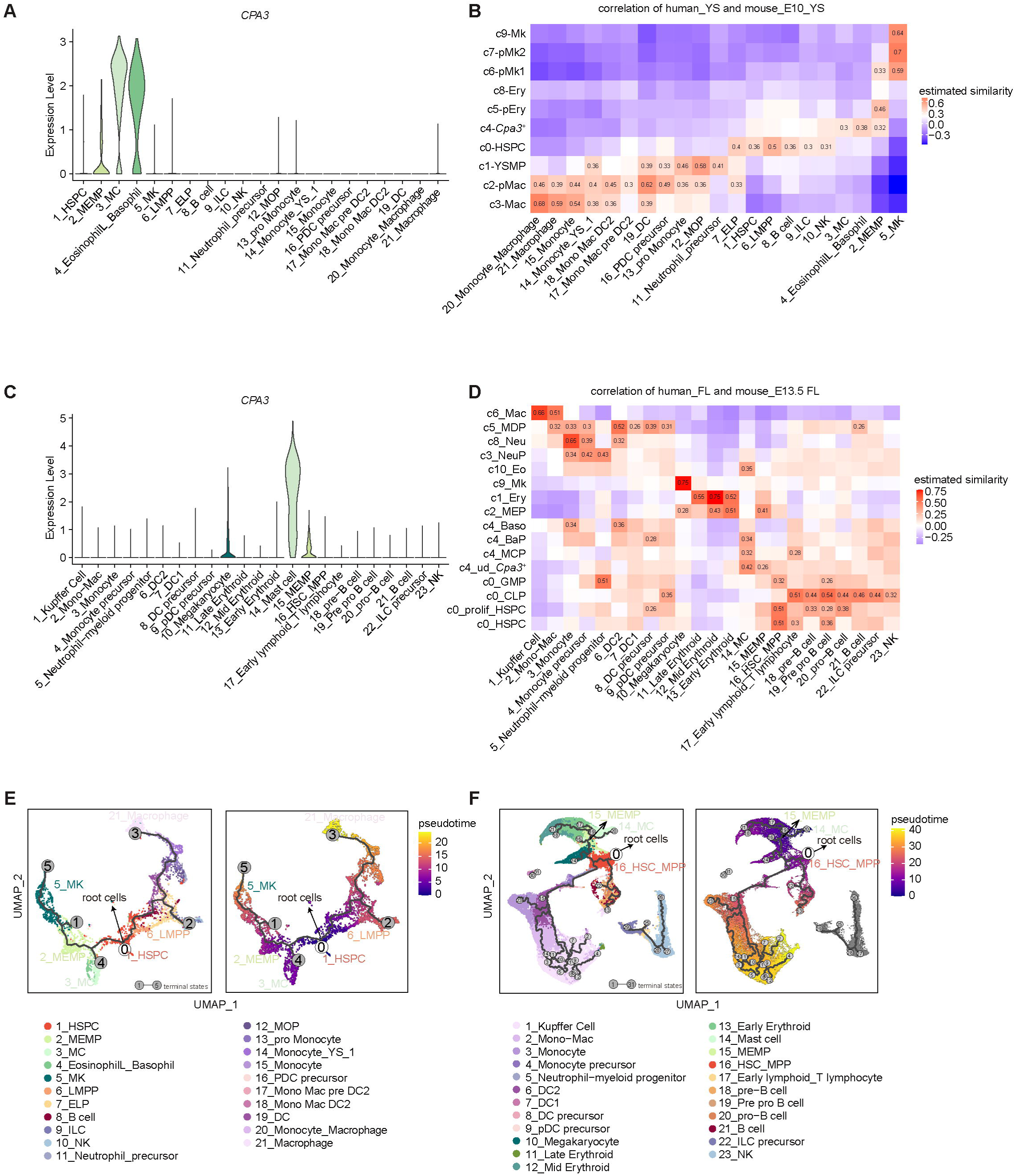
MC development is conserved across species. (A) Violin plot showing the expression of *CPA3* in the specified human YS cell populations (scRNA-seq data from (Popescu et al., 2019)). (B) Pearson correlation analysis between human and murine YS populations (human YS dataset from (Popescu et al., 2019)). (C) Violin plot showing the expression of *CPA3* in the specified human FL cell populations (scRNA-seq data from (Goh et al., 2023)). (D) Pearson correlation analysis between murine FL clusters and human FL populations (human dataset from (Goh et al., 2023)). (E, F) Inferred pseudotime trajectories of cell development in the human YS (E) and FL (F) by monocle3. Lines indicate inferred developmental trajectories. Cells are colored according to identity clusters. UMAP in the right-bottom showing the inferred pseudotime of cells.

## Discussion

In this study, we identified a *Cpa3*-expressing population in E10.0 YS by scRNA-seq, and with genetic fate mapping models we reveal that this *Cpa3*-expressing population sequentially appeared in YS, FL and embryonic skin from E10.5-E14.5, where they could give rise to MCs within a temporal window that is mostly complete around E14.5. And we identified MCPs as Ter119^−^F4/80^−^CD45^+^CD117^+^CD16/32^+^CD135^−^CD115^−^Ly6C^−^CD34^lo^ in FL. Importantly, we found that the embryonic skin MCs are generated by *Cpa3^+^*MCPs but not *Ms4a3*^+^/*Elane*^+^ GMPs. We further showed that human YS and FL contain similar MCP population that corresponds to murine MCPs suggesting a conserved MC developmental program across species.

Unlike hematopoiesis in adult BM, which is relatively stable, embryonic hematopoiesis evolve with the development of the embryo, with different lineages emerge sequentially. While only macrophage, erythrocyte/megakaryocyte/MC lineages could be detected in E10.0 YS, most cell lineages in adult BM could be detected in E13.5 FL, facilitating the transfer of our knowledge in adult BM to embryonic hematopoiesis in FL. Of note, although similar lineages with BM could be found in FL, the phenotype of FL population might differ from their counterpart in BM, for example, E-cadherin marks basophil and MC progenitor in adult BM (Wanet et al., 2021), but only expressed in basophils in FL, emphasizing the need of specific markers for progenitors in FL. Another difference between embryonic and adult BM hematopoiesis is that open niches created by embryo development enables progenitors to give rise to long-lived tissue resident cells, including tissue resident macrophages and MCs. Trajectory analysis suggested that MCs and basophils separate from other myeloid lineages at an early developmental stage. MCPs show a distinct transcriptional profile from GMPs, and GMP fate mapping (*Ms4a3^Cre^* and *Elane^Cre^*) does not label MCPs or MCs, confirming that MCPs arise from earlier progenitors rather than from GMPs. This is in agreement with the result from scRNA-seq data on adult BM, that MCPs arise from early progenitor independent of GMPs (Dahlin et al., 2018), also in agreement with fate mapping result that MMCs arise from Ms4a3^−^ non-GMP BM-derived progenitors (Tauber et al., 2023).

In *Ms4a3^Cre^-Rosa^tdTomato^* mice, basophils were highly labeled while MCs were not (**Figure S4C-D**), suggesting that there may be two separate lineages for generating basophils at an early stage in the FL. In line with this, MCPs showed more similarity to MEPs than to GMPs, accordingly, a population showed both MC and erythrocyte signatures were termed MEMPs in a recent study in humans (Popescu et al., 2019). By comparing the MCPs in FL and the MCs in the body in E13.5 scRNA-seq, we found that the fetal MCs seeded in skin exhibited allergy related membrane surface or secreted proteins similar to adult MCs, including *Tpsg1*, *Tpsb2*, *Fcer1g*, and *Cd9*, indicating that fetal MCs gradually acquired function in the mid embryonic stage, in agreement with a previous report that maternal allergies can be transmitted vertically to the fetus transiently (Msallam et al., 2020).

Another open question regarding MC development is the time window for their generation. CTMCs and most tissue resident macrophages are generated during embryogenesis, and YS macrophage progenitors have been shown to traffic to tissues during a defined developmental window, which is earlier than E14.5 (Stremmel et al., 2018). Our fate mapping experiment with a tamoxifen-inducible Cre (CreERT2) driven by *Zbtb16*, a gene that is expressed by MCPs but not mature MCs, revealed that most MCs were also generated during a similar developmental window with embryonic macrophages. Pulsing *Zbtb16^CreERT2^-Rosa^tdTomato^*before E11.5 labeled MCPs and MCs efficiently, while pulsing at E14.5 did not. This is unlikely to indicate a halt in MCP generation, as MC precursor could still be detected at E15.5 in the blood (Rodewald et al., 1996); rather, it might suggest that the MC “niches” in skin are filled at around E14.5, after which MCs could be largely maintained by self-proliferation, and minor contribution from BM cells after birth. Consistent with this, a large proportion of dermal MCs are proliferating, as shown in Figure 5A. The developmental kinetics of MCs in tissues other than skin needs further investigation, and may vary between tissues (Chia et al., 2023). Furthermore, our results identifying MCPs in mice in agreement with previous reports in humans (Goh et al., 2023; Popescu et al., 2019). Human scRNA-seq data also revealed corresponding *CPA3*^+^ cells in the YS and FL, suggesting that the development of these cells is conserved, and facilitating the transfer of knowledge from studies in mice to the understanding of human MC development.

In conclusion, our study demonstrates that dermal MCs were generated in embryos by *Cpa3^+^* MCPs, independent from GMPs, which seeded in skin largely before E14.5 in the embryonic stage. A clear understanding of the timing and developmental pathway for embryonic MCs may facilitate the design of new approaches to alter MC development or function in order to treat disease or promote health.

## ACKNOWLEDGEMENTS

This work was supported by the National Key R&D Program of China (2021YFA1301400, to B.S.), National Natural Science Foundation of China (NSFC) (31930035 to B.S., 32070880, 32270916, 31900630 to Z.L., 32170895 to N.W.), Research Fund for Foreign Senior Scholars (3231101303 to B.S.), Shanghai Science and Technology Commission(22JC1402600 to B.S.), Shanghai Jiao Tong University Scientific and Technological Innovation Funds (21X010200648 to Z.L.), Shanghai Jiao Tong University 2030 Initiative (WH510363001-16 to Z.L.), Shanghai Rising-Star Program (20QA1408500 to Z.L.). We thank the flow cytometry team, sequencing core, imaging core in Shanghai Institute of Immunology and the Core Facility of Basic Medical Sciences, Shanghai Jiao Tong University School of Medicine for their support. We thank Dr. Lai Guan Ng from Shanghai Immune Therapy Institute, Renji Hospital, Shanghai Jiao Tong University School of Medicine, for providing suggestions on experiments. We thank Dr. Rebecca Gentek from University of Edinburgh, for providing useful comments. We thank Dr. Daniel Ackerman of Insight Editing London for editing this manuscript during preparation.

## AUTHOR CONTRIBUTIONS

W.M., Fei G., H.Z., N.W., S.Z., Y.Z., and Z.X., performed experiments. Z.L. and H.C. analyzed data. Y.Y. provide bioinformatics technical support. Y.L. and B.L. provided material. Z.L., Florent G., and B.S. supervised the study and wrote the manuscript.

## Supplemental Figure Legends

**Figure S1 Single-cell RNA sequencing identifies *Cpa3*-expressing cells in E10.0 YS**

(A-B) *Cpa3* expression in ImmGen (A) and BioGPS (B) databases.

**Figure S2 Fate mapping with *Cpa3^CreERT2^* model reveals that *Cpa3*^+^ progenitors give rise to MCs**

(A) Schematic showing the construction of the *Cpa3^CreERT2^* mouse model. The stop codon of the *Cpa3* gene was replaced with a *2A-CreERT2-pA* cassette.

(B) Representative microscopy images showing the tdTomato^+^ cells (red) in E10.5 *Cpa3^CreERT2^-Rosa ^tdTomato^* embryos pulsed with tamoxifen at E9.5.

(C) Representative microscopy images showing the tdTomato^+^ cells (red) in E16.5 *Cpa3^CreERT2^-Rosa ^tdTomato^* embryos pulsed with tamoxifen at E10.5.

(D) Representative confocal immunofluorescence microscopy images showing the 2 mm back skin of E14.5 and E16.5 *Cpa3^CreERT2^-Rosa ^tdTomato^* embryos pulsed with tamoxifen at E10.5, that were used for cell counting in Figure 2H.

(E) Four thousand sort-purified tdTomato^+^CD45^+^ cells from E13.5 *Cpa3^CreERT2^-Rosa^tdTomato^* embryos that pulsed with tamoxifen at E11.5, were intraperitoneally transferred into Sash (*Kit^W-sh/W-sh^*) mice. tdTomato^+^ cells in the peritoneal cavity were analyzed 2 weeks after transfer.

**Figure S3 Single-cell RNA sequencing identifies MCPs in the FL**

(A) UMAP plot showing the expression of *Cpa3* in FL tdTomato^+^ scRNA-seq in Figure 3A.

(B, C) UMAP plot (B) showing the heterogeneous subclusters within the c0_progenitor cluster. Dot plot (C) showing the expression of top 10 DEGs across subclusters. Colors indicate the average expression of each gene. Spot sizes represent the proportion of gene-expressing cells.

(D) UMAP plot showing the subclusters of FL scRNA-seq data in Figure 3A.

(E, F) Feature plots showing the expression of *Itgb7* (E, left), *Il1rl1* (E, right), *and Cdh1* (F)in *Cpa3^CreERT2^-Rosa^tdTomato^*^+^ FL scRNA-seq data.

(G) Violin plots showing the expression of marker genes used for flow cytometry gating strategy across subclusters in FL.

(H) Feature plot showing the expression of *Cd34* in FL merged scRNA-seq data.

(I) Flow cytometric analysis of tdTomato^+^ cells in the FL of E13.5 *Cpa3^CreERT2^-Rosa^tdTomato^* embryos pulsed with tamoxifen at E11.5.

**Figure S4 MCs do not arise from *Ms4a3*^+^ or *Elane*^+^ GMPs**

(A, B) Feature plots showing the expression of *Ms4a3* (A) and *Elane* (B) in FL *Cpa3^+^*and early progenitor clusters.

(C) Percentage of tdTomato-expressing cells in indicated populations in blood, dermis and peritoneal cavity of adult *Ms4a3^Cre^-Rosa^tdTomato^*mice (n = 3).

(D) Percentage of tdTomato-expressing cells in indicated populations in brain and skin of E14.5 *Ms4a3^Cre^-Rosa^tdTomato^*embryos (n= 5).

(E) Schematic showing the construction of the *Elane^Cre^* model. A *Cre* cassette was inserted at the start codon of *Elane* gene.

**Figure S5 The seeding of MCPs to embryonic skin slows down after E14.5**

(A) Feature plot showing the expression of *Cpa3* in body scRNA-seq data.

(B) The differentially expressed and commonly expressed genes between MCPs in the FL and MCs in skin, including genes encode transcription factors, secreted proteins, and cell membrane proteins.

(C) Heatmap showing the expression of pseudotime-dependent genes presented in Figure S5D.

(D) Schematic showing the construction of the *Zbtb16^CreERT2^* model. A *CreERT2-pA* cassette was inserted at the start codon of the *Zbtb16* gene.

**Figure S6 MC development is conserved across species**

(A) UMAP plot showing the indicated populations in human YS (left). Feature plot showing the expression of *CPA3* (right).

(B) Violin plots showing the expression of *CMA1* across the indicated populations in human YS.

(C) UMAP plot showing the indicated populations in human FL (left). Feature plot showing the expression of *CPA3* (right).

(D) Violin plots showing the expression of *CMA1* across the indicated populations in human FL.

## METHODS

### Lead Contact

Further information and requests for resources and reagents should be directed to and will be fulfilled by the lead contact, Bing Su.

### Materials Availability

Mouse lines generated in this study will be made available upon request. An agreement with our institute’s Materials Transfer Agreement (MTA) may be required.

### Data and Code Availability

The single-cell RNA-seq data will be deposited in public database before publication. This paper does not report original code.

## EXPERIMENTAL MODEL AND SUBJECT DETAILS

### Animals

The following reporter mice have been previously described: B6 ACTb-EGFP (RRID:IMSR JAX:003291) (Okabe et al., 1997), *Ms4a3^Cre^* (RRID:IMSR_JAX:036382) (Liu et al., 2019), and *Rosa26^tdTomato^*(RRID: IMSR_JAX:007914) (Madisen et al., 2010), sash mice (*Kit^W-sh/W-sh^*) (RRID:IMSR_JAX:030764) (Cable et al., 1995; Duttlinger et al., 1995). *Cpa3^CreERT2^*mice (NM-KI-200005) were obtained from Shanghai Model Organisms Center, Inc. *Elane^Cre^* (T006195) were obtained from GemPharmatech Co., Ltd.

*Zbtb16^CreERT2^* mice were generated at the Shanghai Model Organisms Center, Inc. A *CreERT2* cassette was inserted after the start codon of the *Zbtb16* gene via CRISPR-Cas9 in C57BL/6 zygotes. The *Zbtb16^CreERT2^* strain was genotyped by PCR using the following primers:

Wild-type Forward primer 5’-CTTCTCCAGTCCCCTCTGCT-3’

Wild-type Reverse primer 5’-ATGACCACATCGCACAAAGT-3’

Mutant Forward primer 5’-CTTCTCCAGTCCCCTCTGCT-3’

Mutant Reverse primer 5’-CATTGCTGTCACTTGGTCGT-3’ The wild-type band is 211 bp, while the mutant band is 441 bp.

All mice were kept in a specific pathogen-free (SPF) animal facility at the Shanghai Jiao Tong University School of Medicine. All animal experiments were approved by the Institutional Animal Care and Use Committee (IACUC) of Shanghai Jiao Tong University School of Medicine (A2019-042) and were performed in accordance with the University’s guidelines for the care and use of laboratory animals.

## METHOD DETAILS

### Time mating and tamoxifen administration

WT females aged > 8 weeks were time mated with males of indicated genotypes. The presence of vaginal plugs was considered to mark day 0.5 of embryonic development (E0.5). Tamoxifen was prepared by dissolving in corn oil to a final concentration of 17.5 mg/mL and was stored at −20°C. Pregnant mice were given 100 μL of tamoxifen (1.75 mg) by gavage at indicated timepoints.

### Tissue processing and flow cytometry

Tissues from embryos before E14.5 were mechanically dissociated by pipetting and passed through a 70 μm cell strainer to produce a single-cell suspension. For embryos later than E14.5, tissues were cut into small pieces and digested with 0.2 mg/mL collagenase type IV (C5138, Sigma) and 0.05 mg/mL DNase I (Cat# 1 0104159 001, Roche) in RPMI 1640 medium with 10% FCS at 37°C for 30 min. Cells were then dissociated by pipetting and passing through a 70 μm cell strainer. For immune staining, nonspecific antibody binding to cells was blocked by incubation with a purified anti-CD16/32 antibody (clone 2G8; BD Biosciences) at 4°C for 15 min, and the cells were stained with fluorophore-conjugated antibodies and/or biotin-conjugated antibodies at 4°C for 25 min. Cells were washed and further stained with fluorophore-conjugated streptavidin to detect biotinylated CD135 antibody. Cells were stained with DAPI and filtered with a 70 μm cell strainer before data acquisition on a BD Fortessa X20 or Symphony flow cytometer (BD Biosciences). Data were analyzed with FlowJo (FlowJo LLC).

### Giemsa and toluidine blue staining

Sort-purified cells were spun onto glass slides using a Cytospin 4 Cytocentrifuge (Thermo Scientific), then air-dried at room temperature for 20 min. For Giemsa staining, slides were stained with Wright-Giemsa solution A for 1 min and Wright-Giemsa solution B for 10 min. For toluidine blue staining, slides were stained with toluidine blue solution for 30 min. After staining, slides were washed with flow water, air-dried and sealed with mounting medium. Images were acquired with an Olympus BX53 light microscope equipped with a 100× oil immersion objective lens.

### Immunofluorescence

Embryos were fixed overnight in fixation buffer containing 2% paraformaldehyde and dehydrated in 30% sucrose before embedding in OCT freezing media (Sakura). Sections were cut to 20 μm on a Leica cryostat. For immunofluorescence, sections were blocked for 1 h at room temperature in blocking buffer containing 1% normal mouse serum, 1% BSA and 0.3% Triton X-100. The sections were stained with the indicated fluorophore-conjugated antibodies overnight at 4°C in a dark, humidified chamber. Antibodies for immunofluorescence can be found in the Key Resources Table. Images were captured under an Olympus FV3000 confocal microscope.

### Adoptive cell transfer

Four thousand sort-purified tdTomato^+^CD45^+^ cells from E13.5 *Cpa3^CreERT2^-Rosa^tdTomato^* embryos FL that pulsed at E11.5, were intraperitoneally transferred into Sash (*Kit^W-sh/W-sh^*) mice. tdTomato^+^ cells in the peritoneal cavity were analyzed 2 weeks after transfer.

### Cell Culture

Sort-purified tdTomato*^+^* MCPs from the FL of E13.5 *Cpa3^CreERT2^-Rosa^tdTomato^* embryos pulsed with tamoxifen at E11.5, were cultured in medium (RPMI 1640 with 10% FCS and 1% penicillin-streptomycin) supplemented with 10 ng/ml IL-3 (PeproTech) and 10 ng/ml SCF (PeproTech) *in vitro* (Li et al., 2018; Wang et al., 2014). Cells were analyzed by flow cytometry after 4 days.

### Single-cell RNA sequencing

For scRNA-seq of YS, WT female mice were crossed with an B6 ACTb-EGFP male. YS of E10.0 embryos were harvested and digested with 0.2 mg/mL collagenase type IV (C5138, Sigma) and 0.05 mg/mL DNase I (Roche) in RPMI 1640 medium with 10% FCS at 37°C for 30 min and dissociated by pipetting. Cell suspensions were stained and FACS-sorted for both CD45^+^ and CD41^+^ cells. For scRNA-seq of FL and body wall, WT female mice were crossed with a *Cpa3^CreERT2^-Rosa^tdTomato^* male. Pregnant females were pulsed with one dose of tamoxifen at E11.5, and FLs and body walls were harvested at E13.5. Tissues were dissociated by pipetting. Tissue cell suspensions were stained and FACS-sorted for tdTomato^+^ and tdTomato^−^CD45^+^ cells, respectively. Sort-purified cells were resuspended in PBS with 0.04% bovine serum albumin (BSA) to a concentration of around 1,000 cells per μL. Cells were loaded on a Chromium Single Cell Controller (10x Genomics). Single-cell transcriptomics were generated according to the manufacturer’s instructions. Sequencing was performed on an Illumina NovaSeq 6000 platform.

### Data alignment and preprocessing for scRNA-seq

Raw reads from scRNA-seq were processed through the Cell Ranger pipeline (version 4.0.0, 10x Genomics) with alignment to the mm10 mouse genome, generating gene-cell barcode matrices. The matrices underwent further refinement with the Seurat package (version 3.2.3), filtering out cells and genes based on a defined set of parameters to refine the dataset for further analysis. Cells with ≥ 200 genes and genes expressed in ≥ 3 cells were filtered according to certain conditions (nFeature_RNA > 200 & nFeature_RNA < 7500 & percent.mito > −Inf & percent.mito < 0.05 & percent.Hbb-bs < 10 & nCount_RNA > 800 & nCount_RNA < 10000) for downstream analysis.

### Cell clustering and dimensionality reduction

After filtering, the data were normalized and scaled by regressing the percentage of mitochondrial genes to mitigate their impact on the dataset’s variance. Identification of the 2000 most variable genes was performed via the “FindVariableFeatures” function in Seurat, setting the stage for Principal Component Analysis (PCA) to reduce dataset dimensionality. Graph-based clustering was achieved using “FindNeighbors”, while “FindClusters” was utilized to delineate cell subtypes. Subtype visualization was accomplished using the UMAP algorithm, with differential expression of marker genes assessed via the Wilcoxon rank sum test, employing “FindAllMarkers” with predefined thresholds (min.pct and logfc.threshold parameters set to 0.25).

### Transcriptional dynamics analysis

Transcriptional trajectory analysis was conducted using Monocle3 (version 1.0.0), converting the Seurat dataset into a Monocle3-compatible format through SeuratWrappers (version 0.3.1). The trajectory partitioning and cell clustering were executed via the “cluster_cells” function, which were then integrated with the Seurat-derived clusters. The trajectory construction was based on the “learn_graph” function, and pseudotime ordering of cells was determined using “order_cells”, with the initiation points manually assigned for each cell type. Spatial autocorrelation in gene expression across multiple dimensions was quantified using Moran’s *I* statistic within the “graph_test” function, retaining only genes with high-confidence results to define gene expression modules, which were subsequently identified using “find_gene_modules” at a specified resolution.

### Similarities in alignment between humans and mice

1. *Sample collection and processing.* Human YS and liver single-cell RNA sequencing datasets were curated from publicly available resources (Goh et al., 2023; Villar and Segura, 2020), ensuring the data adhered to the respective repositories’ ethical and quality standards. The criteria used to select these datasets required the CD45^+^ cells to be isolated and a minimum threshold of 500 cells per sample to guarantee dependable statistical analysis.
2. *Gene expression analysis*. For each cell subtype within the human and murine datasets, the average gene expression value was calculated to ascertain the representative expression profile.
3. *Quantile normalization and batch effect mitigation.* Prior to comparative analysis, the human and murine scRNA-seq data were subjected to batch effect mitigation to correct for technical variations inherent to sample processing and sequencing runs. This was achieved by implementing quantile normalization across the datasets to standardize the distribution of expression values. Thereafter, the ComBat algorithm, a widely used tool for adjusting batch effects in genomic data, was employed to harmonize the datasets, ensuring that biological conclusions were not confounded by technical artifacts.
4. *Subgroup similarity assessment by Pearson correlation analysis.* To analyze the similarity between cell subgroups across human and mouse datasets, high-variance genes and top30 marker genes were intersected to create a focused set for comparison. The Pearson correlation coefficient was computed for each gene across species within the same cell subtypes to determine the degree of expression similarity. This cross-species comparative analysis aimed to elucidate conserved patterns of gene expression that may underpin common functional characteristics across human and mouse immune cell lineages.

## QUANTIFICATION AND STATISTICAL ANALYSIS

### Statistical analysis

Statistical analyses were performed using Prism software (GraphPad). Two-tailed Student’s *t*-test was used when comparing two groups. *p* < 0.05 was considered significant: \**p* < 0.05; \*\**p* < 0.01; \*\*\**p* < 0.001. *p* > 0.05 was considered nonsignificant (n.s.). Number of replicates, sample size, and significance tests for each experiment are specified in the Figure legends.

