## Supplementary figures and images for "Embryonic Mast Cells Arise from *Cpa3*-expressing Precursors Independent of Granulocyte-Monocyte Progenitors"

### Supplemental Figure 1

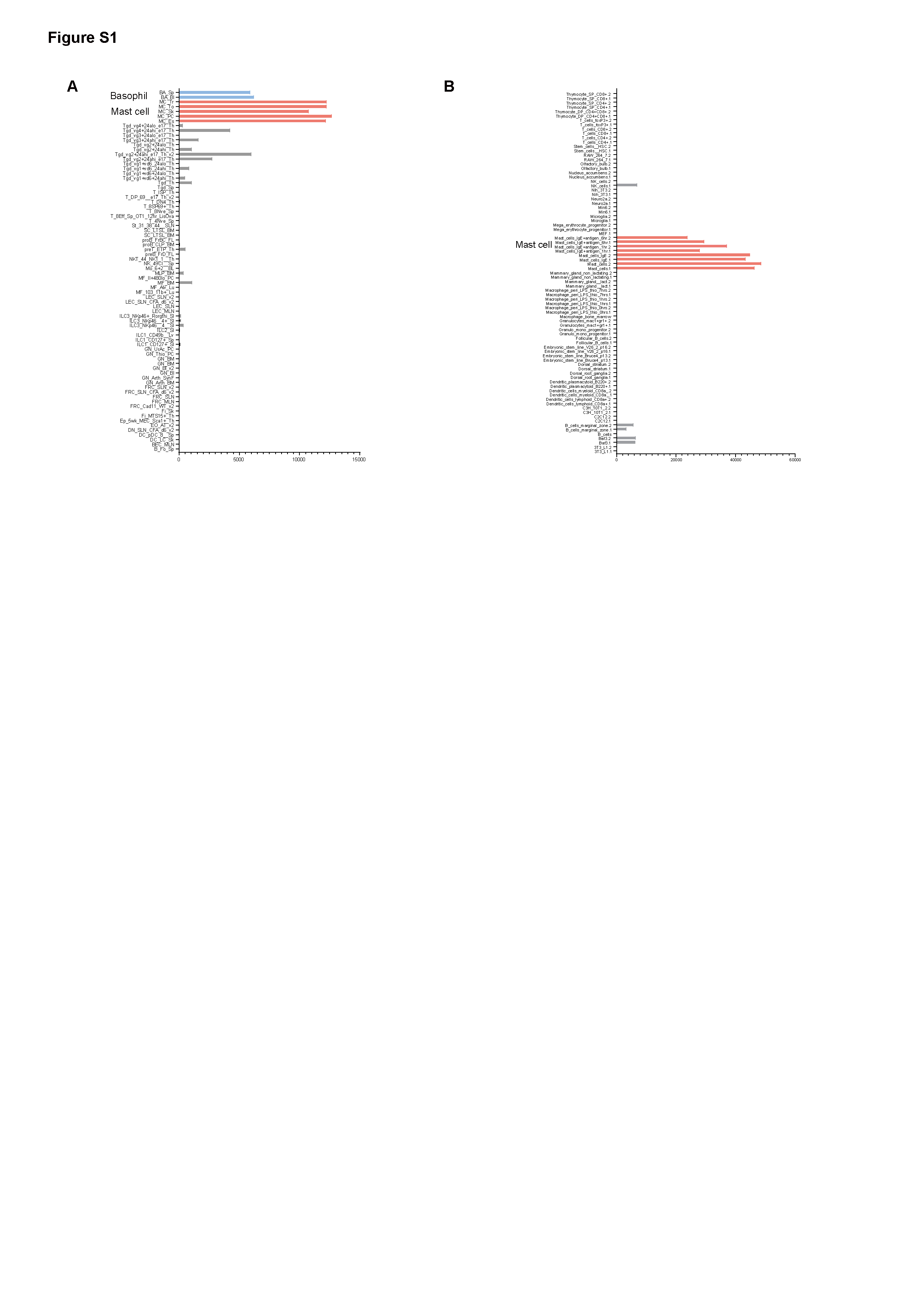

### Supplemental Figure 2

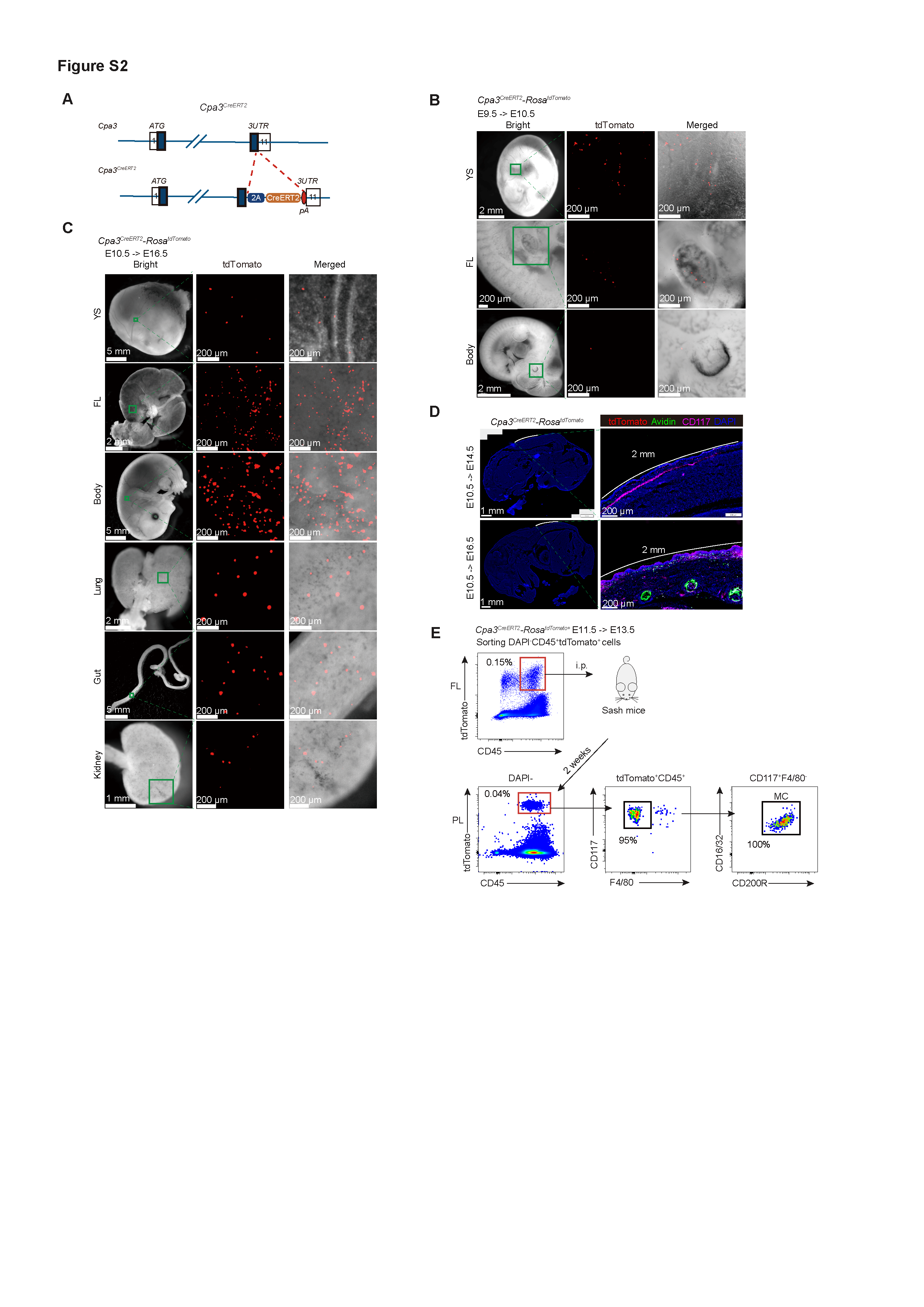

### Supplemental Figure 3

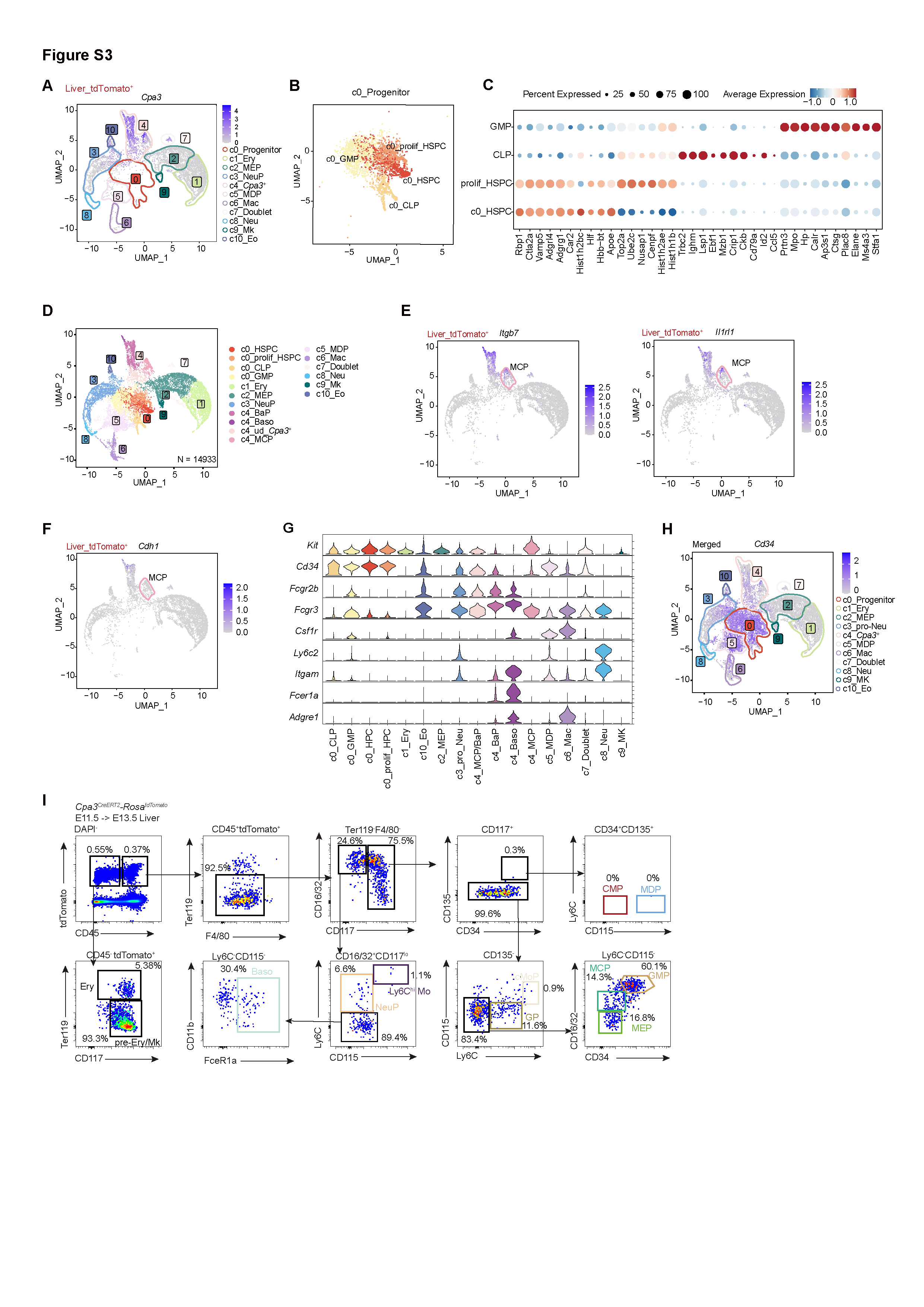

### Supplemental Figure 4

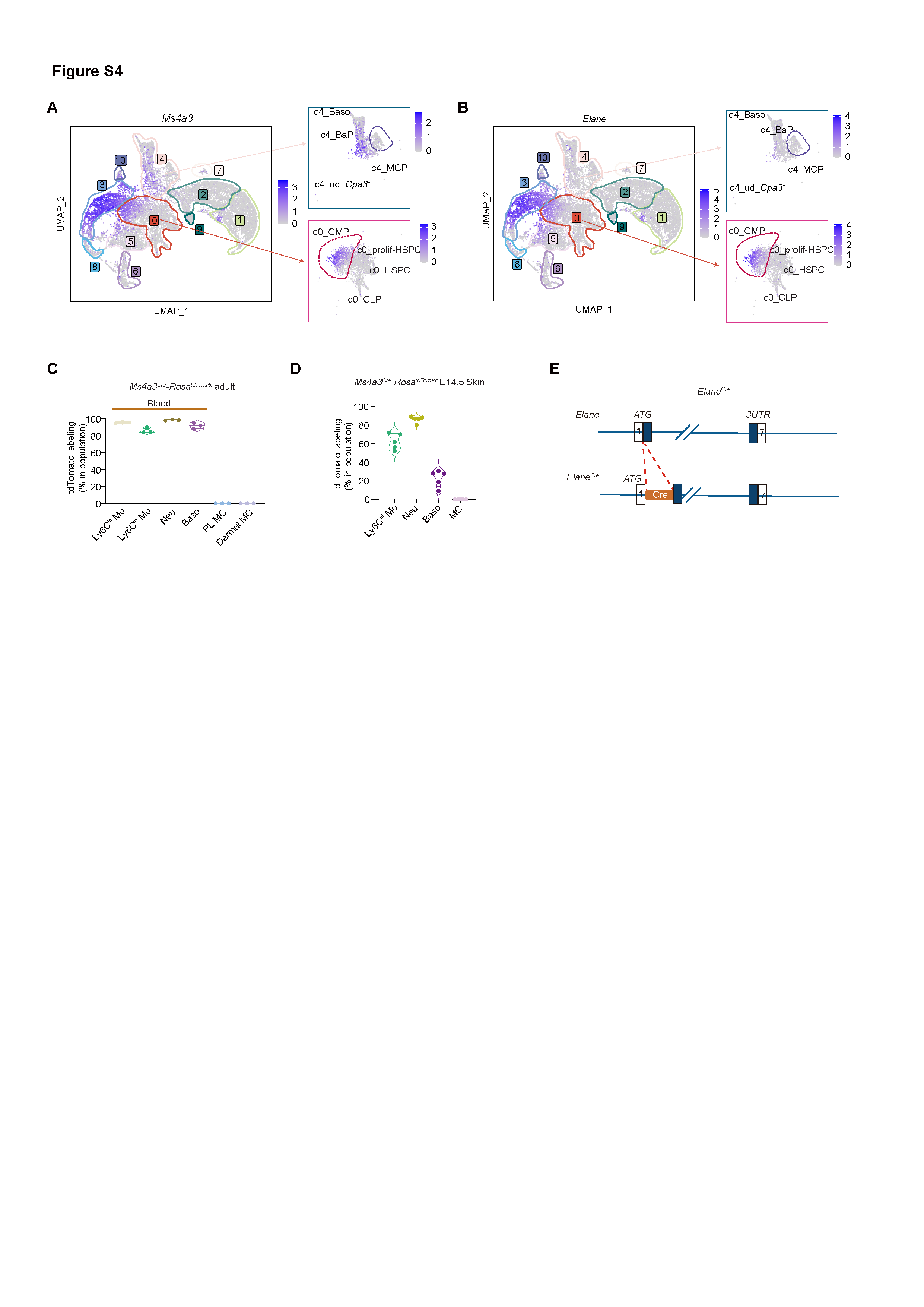

### Supplemental Figure 5

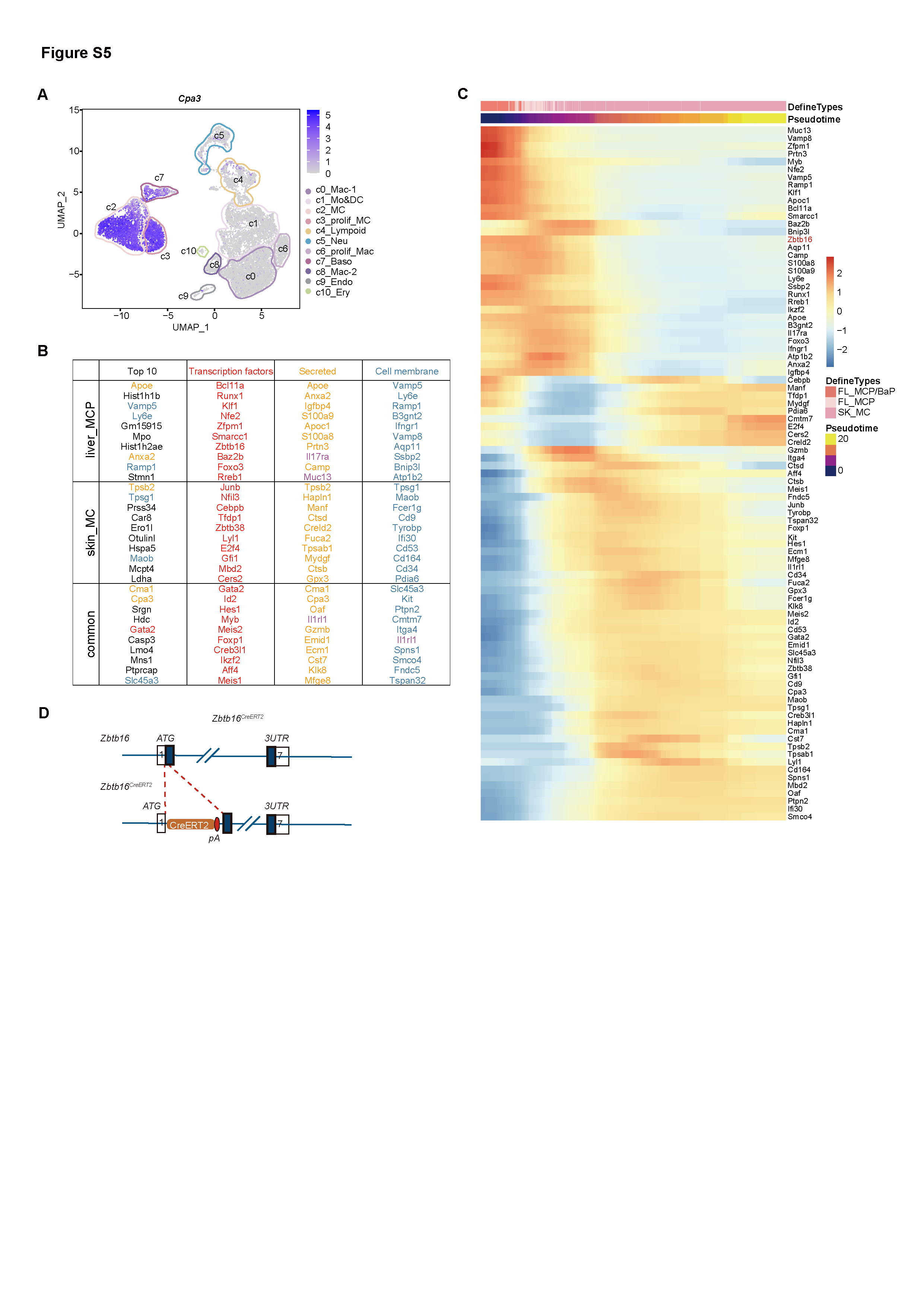

### Supplemental Figure 6

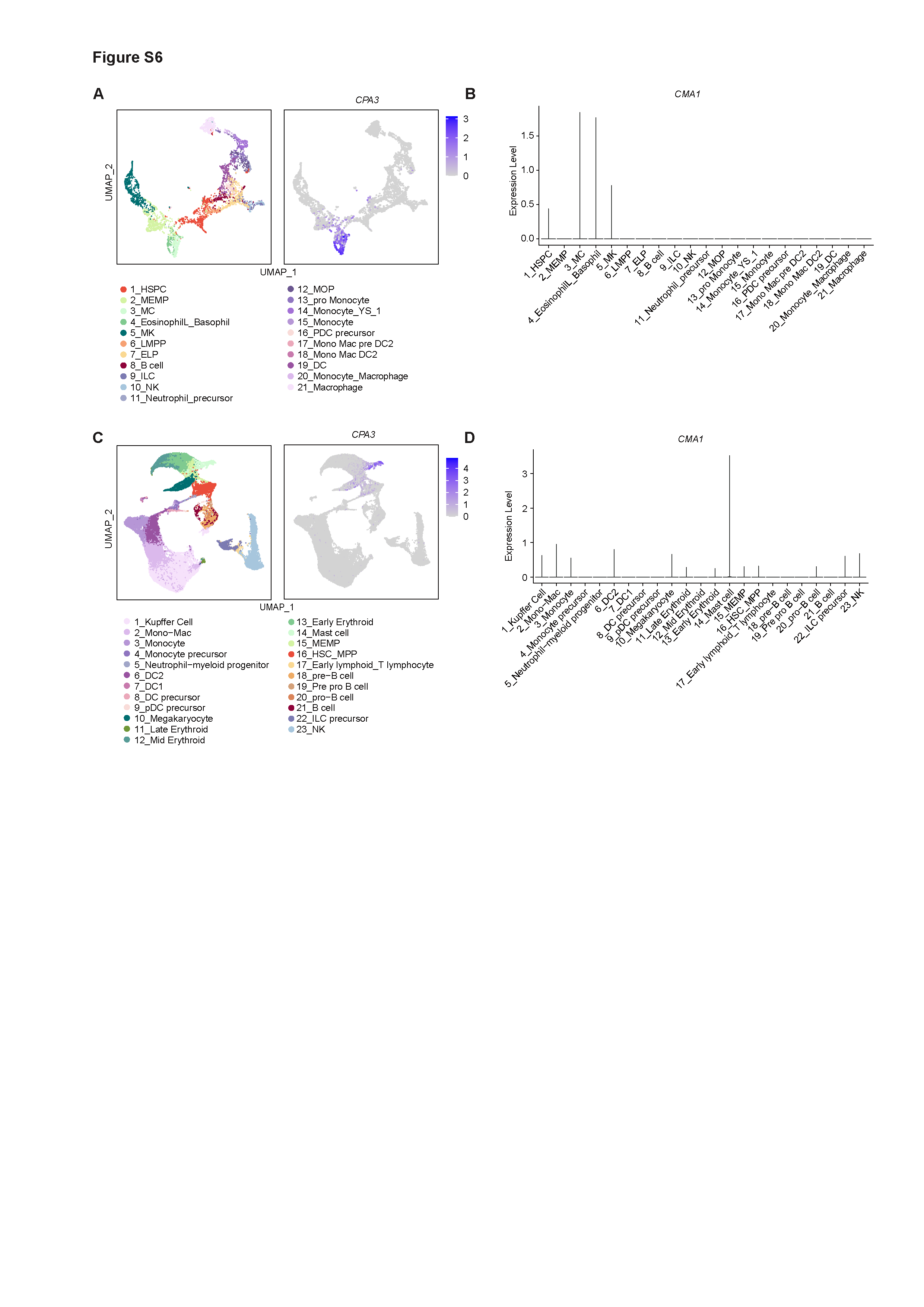
