## Supplementary material for "Embryonic Mast Cells Arise from *Cpa3*-expressing Precursors Independent of Granulocyte-Monocyte Progenitors": material

| REAGENT or RESOURCE | SOURCE | IDENTIFIER |
|---|---|---|
| Antibodies | | |
| Anti-mouse Sca-1/Ly6A Alexa Fluor 700 (D7) | eBioscience | Cat#56-5981-82; RRID: AB_657836 |
| Anti-mouse MHCII Alexa Fluor 700 (M5/114.15.2) | eBioscience | Cat#56-5321-82; RRID: AB_494009 |
| Anti-mouse Ter-119 APC (TER-119) | Biolegend | Cat#116212; RRID: AB_313713 |
| Anti-mouse CD3e APC (145-2C11 ) | Biolegend | Cat#100312; RRID: AB_312677 |
| Anti-mouse CD19 APC (1D3) | eBioscience | Cat#17-0193-82; RRID: AB_1659676 |
| Anti-mouse NK1.1 APC (PK136) | Biolegend | Cat#108709; RRID: AB_313396 |
| Anti-mouse Ly6g BV711 (1A8) | Biolegend | Cat#127643; RRID: AB_2565971 |
| Anti-mouse Siglec F AF647 (E50-2440) | BD | Cat#562680; RRID: AB_2687570 |
| Anti-mouse CD200R APC (OX110) | eBioscience | Cat#17-5201-82; RRID: AB_10717289 |
| Anti-mouse CD16/32 APC/Cy7 (93) | Biolegend | Cat#101328; RRID: AB_2104158 |
| Anti-mouse CD64 APC (X54-5/7.1) | Biolegend | Cat#139306;RRID:AB_11219391 |
| Anti-mouse CD135 biotin (A2F10) | eBioscience | Cat#13-1351-82; RRID: AB_466599 |
| Anti-mouse CD45 BUV395 (104) | BD | Cat#564279; RRID: AB_2651134 |
| BUV737 Streptavidin | BD | Cat#612775; RRID: AB_2869560 |
| Anti-mouse CD34 BV421 (SA376A4) | Biolegend | Cat#152208; RRID: AB_2650766 |
| Anti-mouse Siglec F BV421 (E50-2440) | BD | Cat#562681; RRID: AB_2722581 |
| Anti-mouse CD117 cKit BV510 (2B8) | Biolegend | Cat#105839; RRID: AB_2629798 |
| Anti-mouse CD115 BV605 (AFS98) | Biolegend | Cat#135517; RRID: AB_2562760 |
| Anti-mouse CD206 BV605 (C068C2) | Biolegend | Cat#141721; RRID: AB_2562340 |
| Anti-mouse CD11b BV650 (M1/70) | Biolegend | Cat#101259; RRID: AB_2566568 |
| Anti-mouse Ly6g BV711 (1A8) | Biolegend | Cat#127643; RRID: AB_2565971 |
| Anti-mouse Ly6c BV785 (HK1.4) | Biolegend | Cat#128041; RRID: AB_2565852 |
| Anti-mouse F4/80 AF488 (BM8) | Biolegend | Cat#123120; RRID: AB_893479 |
| Anti-mouse FceRI PE-Cy7 (MAR-1) | eBioscience | Cat#25-5898-80; RRID: AB_2573492 |
| Anti-mouse CD172a PerCP-eFluor® 710 (P84) | eBioscience | Cat#46-1721-82; RRID: AB_10804639 |
| Anti-mouse CD117 AF647 (2B8) | Biolegend | Cat#105815 ;RRID:AB_493473 |
| Anti-mouse Avidin AF488 (BM8) | Biolegend | Cat#123120;RRID:AB_893491 |
| Anti-mouse F4/80 PE (BM8) | Biolegend | Cat#123109;RRID:AB_893498 |
| Chemicals, Peptides, and Recombinant Proteins | | |
| Collagenase type IV | Sigma | Cat# C5138 |
| DNase I | Roche | Cat# 1 0104159 001 |
| Dispase | Gibco | Cat# 17105-041 |
| Tamoxifen | Sigma | Cat# T5648 |
| DAPI | ThermoFisher | Cat# D1306 |
| Corn oil | Aladdin | C116025-500ml |
| Proteinase K | Yeasen | 10401ES80 |
| 2 × Rapid Taq Master Mix | Vazyme | P222-03 |
| 2000 DNA Marker | Yeasen | 10501ES60 |
| 50xTAE | Beyotime | 31-008 |
| DEPC water | Beyotime | R0022 |
| Triton X-100 | Beyotime | A620554 |
| Tris HCL | Sigma Aldrich | PHR9293 |
| Dispase | Gibco | 17105-041 |
| ImmEdge Pen | Vector | H-4000 |
| $NH_4Cl$ | Sigma Aldrich | 901230 |
| BSA | Yeasen | 36101ES25 |
| FBS | Gibco | 10437-028 |
| DMEM, high glucose, pyruvate | Thermofisher | 11995-065 |
| RPMI 1640 Medium | Thermofisher | 11875093 |
| IL-3 | PeproTech | 213-13 |
| SCF | PeproTech | 250-03 |
| $KHCO_3$ | Sigma Aldrich | 237205 |
| Giemsa | Baso | BA4017 |
| Toluidine | Servicebio | G1032-100ML |
| Critical Commercial Assays | | |
| SMART-Seq HT Kit | TaKaRa | Cat# 634437 |
| TruePrep DNA Library Prep Kit V2 for Illumina | Vazyme | Cat# TD 502-02 |
| Chromium Next GEM Single Cell 3′ Reagent Kits | 10x Genomics | v3.1 |
| Deposited Data | | |
| Bulk mRNA-seq data | This paper | |
| Single-cell RNA-seq data | This paper | |
| Experimental Models: Organisms/Strains | | |
| Mouse: *Ms4a3*$^{Cre}$ | Liu et al., 2019 | RRID: IMSR_JAX:036382 |
| Mouse: *Cpa3*$^{CreERT2}$ | Model organism | NM-KI-200005 |
| Mouse: *Zbtb16*$^{CreERT2}$ | Model organism | N1-1725 |
| Mouse: *Elane*$^{Cre}$ | Gempharmatech | T006195 |
| Mouse: C57BL/6-Tg(CAG-EGFP)1Osb/J | The Jackson Laboratory | RRID: IMSR_JAX:003291 |
| Mouse: B6.Cg-KitW-sh/HNihrJaeBsmGlliJ | The Jackson Laboratory | RRID: IMSR_JAX:012861 |
| Mouse: *Rosa26*$^{tdTomato}$ | The Jackson Laboratory | RRID: IMSR_JAX:007914 |
| Oligonucleotides | | |
| Genotyping primer sequences | This paper | See Method Details |
| Software and Algorithms | | |
| FlowJo V10 | FlowJo LLC | https://www.flowjo.com |
| GraphPad Prism 6 | GraphPad Software | https://www.graphpad.com |
| Imaris | Bitplane | http://www.bitplane.com |

| | | |
|---|---|---|
| R v4.1.2 | The Comprehensive R Archive Network | https://cran.r-project.org/ |
| Software and Algorithms | | |
| FlowJo V10 | FlowJo LLC | https://www.flowjo.com |
| GraphPad Prism 6 | GraphPad Software | https://www.graphpad.com |
| Imaris | Bitplane | http://www.bitplane.com |
| R v4.1.2 | The Comprehensive R Archive Network | https://cran.r-project.org/ |
